# Mapping immunodominant sites on the MERS-CoV spike glycoprotein targeted by infection-elicited antibodies in humans

**DOI:** 10.1101/2024.03.31.586409

**Authors:** Amin Addetia, Cameron Stewart, Albert J. Seo, Kaitlin R. Sprouse, Ayed Y Asiri, Maha Al-Mozaini, Ziad A Memish, Abeer Alshukairi, David Veesler

## Abstract

Middle-East respiratory syndrome coronavirus (MERS-CoV) first emerged in 2012 and causes human infections in endemic regions. Most vaccines and therapeutics in development against MERS-CoV focus on the spike (S) glycoprotein to prevent viral entry into target cells. These efforts, however, are limited by a poor understanding of antibody responses elicited by infection along with their durability, fine specificity and contribution of distinct S antigenic sites to neutralization. To address this knowledge gap, we analyzed S-directed binding and neutralizing antibody titers in plasma collected from individuals infected with MERS-CoV in 2017-2019 (prior to the COVID-19 pandemic). We observed that binding and neutralizing antibodies peak 1 to 6 weeks after symptom onset/hospitalization, persist for at least 6 months, and broadly neutralize human and camel MERS-CoV strains. We show that the MERS-CoV S_1_ subunit is immunodominant and that antibodies targeting S_1_, particularly the RBD, account for most plasma neutralizing activity. Antigenic site mapping revealed that polyclonal plasma antibodies frequently target RBD epitopes, particularly a site exposed irrespective of the S trimer conformation, whereas targeting of S_2_ subunit epitopes is rare, similar to SARS-CoV-2. Our data reveal in unprecedented details the humoral immune responses elicited by MERS-CoV infection, which will guide vaccine and therapeutic design.

## Introduction

Coronaviruses (CoV) are zoonotic pathogens causing significant morbidity and mortality in humans as exemplified by the recent emergence of severe acute respiratory syndrome coronavirus (SARS-CoV-1), Middle-East respiratory syndrome coronavirus (MERS-CoV), and severe acute respiratory syndrome coronavirus-2 (SARS-CoV-2) (*1–6*). MERS-CoV first emerged in 2012 and human infections have continued to be documented in endemic regions, including in 2024 (*7, 8*). MERS-CoV causes severe respiratory illness in infected individuals with a reported case fatality rate of approximately 35% (*7, 9*). Sustained human-to-human MERS-CoV transmission infrequently occurs with the majority of new cases occurring through spillover from infected camels to humans (*10, 11*).

MERS-CoV entry into target cells is facilitated by the viral spike (S) glycoprotein (*12, 13*), which consists of two subunits, S_1_ and S_2_ (*14*). The S_1_ subunit contains the receptor-binding domain (RBD) and the N-terminal domain (NTD), that permit attachment to the target cell through interactions with dipeptidyl peptidase 4 (DPP4) and sialosides, respectively (*12, 15–18*). The S_2_ subunit is the machinery promoting fusion of the viral and host membranes to initiate infection (*14, 19*). Viral entry involves proteolytic cleavage of MERS-CoV S by either furin (during viral morphogenesis) or cathepsins (after virions are endocytosed) generating the S_1_ and S_2_ subunits that remain non-covalently linked in the S trimer. Subsequent S_2_ subunit cleavage by TMPRSS2 (at the plasma membrane) or cathepsins (in the endosomes) yields S_2_’ and allows large-scale conformational changes to fuse the viral and host membranes (*13, 19–21*). Similar to SARS-CoV-2, S-directed neutralizing antibodies have been suggested to be the primary correlate of protection against MERS (*22–24*). As such, MERS-CoV vaccines currently under development encode MERS-CoV S and aim to elicit potent neutralizing antibody responses that block viral entry (*22, 25–28*).

Several monoclonal neutralizing antibodies (mAbs) targeting MERS-CoV S have been isolated and characterized to date (*29–43*). They recognize epitopes located in the RBD, NTD, or S_2_ subunit, indicating that each of these MERS-CoV S domains are targets of neutralizing mAbs. Although these results establish proof-of-principle of mAb-mediated viral inhibition, and in vivo protection for a subset of them, they do not provide any information on the quantitative contribution of each S domain or antigenic site to polyclonal plasma binding and neutralizing activity. Studies of antibody responses elicited by SARS-CoV-2 infection and vaccination revealed that the SARS-CoV-2 RBD is immunodominant (*44, 45*) and that antibodies targeting the SARS-CoV-2 RBD account for most plasma neutralizing activity against infection/vaccine-matched and mismatched viruses such as variants (*46–49*). Hence, analysis of MERS-CoV infection-elicited polyclonal antibody responses in humans may provide insights into preferred S domains and epitopes to target for developing MERS-CoV vaccines and therapeutics.

Here, we analyzed binding and neutralizing antibody titers in plasma samples collected from individuals who were hospitalized with MERS-CoV infections prior to the SARS-CoV-2 pandemic. S-directed binding and neutralizing antibody titers peaked rapidly and persisted for at least 6 months after symptom onset or hospitalization. We determined that infection-elicited antibodies broadly neutralize human and camel MERS-CoV variants and that S_1_-directed antibodies, and more specifically RBD-directed antibodies, account for the majority of plasma neutralizing activity. Lastly, we observed that polyclonal antibodies predominantly recognize RBD epitopes whereas the S_2_ subunit is much more rarely targeted. Collectively, our findings reveal the immunodominant sites on the MERS-CoV S protein that contribute to the neutralization potency of infection-elicited plasma antibodies.

## Results

### MERS-CoV infection induces S-directed plasma binding and neutralizing antibodies

To understand humoral immune responses following MERS-CoV exposure, we collected plasma samples from 30 individuals hospitalized with MERS between 2017 and 2019 (i.e. prior to the COVID-19 pandemic). The plasma donors ranged from 21 to 80 (median: 52; **Table S1**) years of age and 27 of the 30 individuals had at least one underlying comorbidity. A total of 98 plasma samples were obtained with each subject contributing 1 to 12 samples collected 3 to 191 days post-symptom onset or hospitalization (**Table S2**).

Analysis of plasma binding antibody titers against the prefusion MERS-CoV S 2P ectodomain trimer (EMC/2012 strain; GenBank accession no. NC_019843.3) by ELISA showed that S-directed IgGs were present in 97 out of 98 plasma samples with half-maximal effective dilution (ED_50_) titers ranging from ≤10 to 55,978 (geometric mean titer, GMT: 875; **Figure 1A**, **Figure S1**). Evaluation of the corresponding neutralizing activity using VSV pseudotyped with the MERS-CoV EMC/2012 S revealed that 95 out of 98 plasma samples had neutralizing antibody titers ranging between half-maximal inhibitory dilutions (ID_50_) of ≤10 and 2,120 (GMT: 192; **Figure 1B**, **Figure S2**). We found that plasma binding antibody titers positively correlated with neutralizing activity (Spearman r = 0.78) when the 97 samples with detectable S binging titers were included in the analysis as well as when only the sample with the highest S IgG binding titer from each individual was included (Spearman r = 0.81; **Figure S3**).

**Figure 1.**
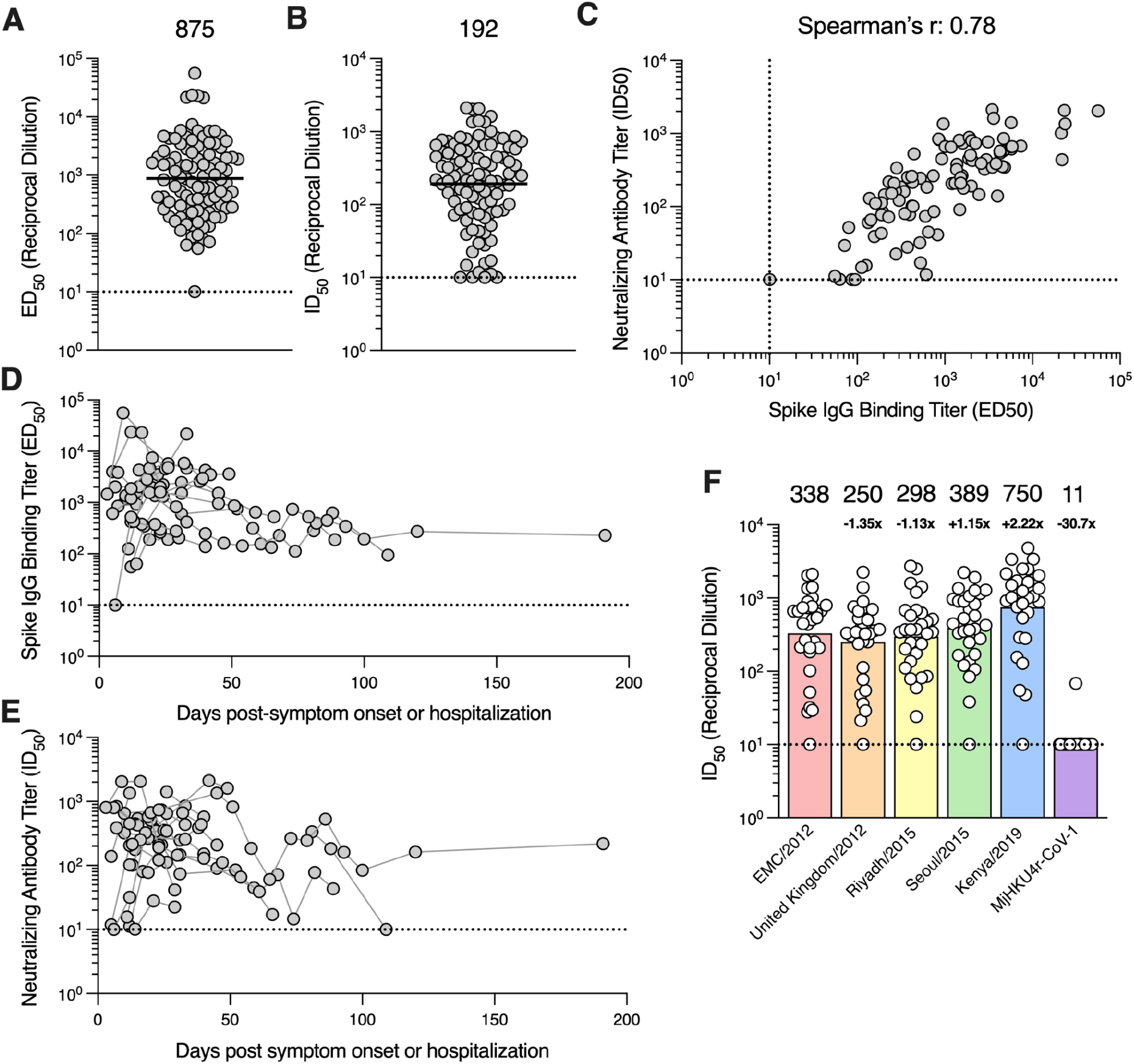
MERS-CoV infection induces robust S binding and neutralizing antibody responses. **A)** IgG binding titers (half-maximal effective dilution [ED_50_]) against the prefusion MERS-CoV EMC/2012 S ectodomain trimer and **B)** neutralizing antibody titers (half-maximal inhibitory dilution [ID_50_]) against VSV pseudotyped with the MERS-CoV EMC/2012 S measured for 98 plasma samples collected from 30 individuals hospitalized with MERS-CoV infection between 2017-2019 in Saudi Arabia. The geometric mean titer (GMT) is represented by the black bar and displayed above the plot and the limit of detection (ED_50_ or ID_50_: 10) is represented by the dashed line. Data reflect results obtained from one biological replicate and are representative of data obtained from at least two biological replicates conducted with unique batches of S protein or pseudovirus. **C)** Correlation analysis between neutralizing antibody and S IgG binding titers in the 97 plasma samples with detectable S binding titers. Kinetics of the **D)** S IgG binding titers and **E)** neutralizing antibody titers for 90 plasma samples collected from 22 individuals who contributed two or more samples. Samples collected from the same individual are connected with gray lines. **F)** Neutralization potency (ID_50_) of plasma samples against VSV pseudotyped with the spike protein of the indicated MERS-CoV variant or the related merbecovirus MjHKU4r-CoV-1. Only the sample with highest S binding titer per individual was included in the analysis. GMTs are represented by the bars and displayed above the plot. The fold-change in GMT compared to MERS-CoV EMC/2012 pseudovirus is indicated below the GMT for each pseudovirus. Data reflect results obtained from one biological replicate and are representative of data obtained from at least two biological replicates conducted with unique batches of pseudoviruses.

### Durability of antibody responses after MERS-CoV infection

To understand the kinetics of elicitation and waning of antibody responses in MERS-CoV plasma, we examined binding and neutralizing antibody titers across 90 plasma samples from 22 individuals who contributed two or more longitudinal samples. Time points were recorded as the number of days post-symptom onset or hospitalization. All 22 individuals exhibited detectable S-directed IgG binding and neutralizing activity 12 days post-symptom onset or hospitalization and some of them as early as 3 days post-symptom onset or hospitalization (**Figure 1D-E**). Peak binding and neutralizing antibody titers occurred 7 to 38 days and 6 to 45 days post-symptom onset/hospitalization, respectively, based on analysis of 21 subjects contributing at least two samples collected in the first 60-day window. Binding and neutralizing antibody titers were detectable for up to 191 days post-symptom onset/hospitalization, with the exception of one individual without plasma neutralizing activity remaining at day 109, consistent with previous reports (*50–54*). These data show that antibody responses peak rapidly following MERS-CoV infection and are durable over time, although we observed some heterogeneity in the cohort evaluated.

### MERS-CoV infection elicits broadly neutralizing antibodies against variants

To understand the breadth of neutralizing antibody responses elicited by infection, we examined plasma neutralization against VSV pseudotyped with a panel of MERS-CoV S variants including United Kingdom/2012 (GenBank accession no. NC_038294.1), Riyadh/2015 (GenBank accession no. KT806049.1), Seoul/2015 (GenBank accession no. KT374056.1), and Kenya/2019 (GenBank accession no. OK094446.1). MERS-CoV Kenya/2019 was identified from a camel during an outbreak in Kenya (*55*) while the other variants were identified from human cases. These variants harbor between 2 and 8 S amino acid substitutions relative to EMC/2012 (Table S3) with United Kingdom/2012 S and Seoul/2015 S containing mutations known to reduce the neutralizing activity of mAbs or sera (*56–58*). The D510G S mutation of the Seoul/2015 strain has further been shown to reduce DPP4 binding affinity and impair viral entry in human cells (*59, 60*).

Relative to the EMC/2012 strain (GMT: 338), plasma neutralizing antibody titers were reduced 1.35-fold against the United Kingdom/2012 S VSV (GMT: 250; **Figure 1E**; **Figure S4**), 1.13-fold against the Riyadh/2015 S VSV (GMT: 298) and slightly increased against Seoul/2015 S VSV (GMT: 389). Strikingly, neutralizing activity was enhanced 2.22-fold against Kenya/2019 S VSV (GMT: 750), suggesting that recent human MERS-CoV infections are likely due to zoonotic spillover from camels. This interpretation is consistent with prior phylodynamic analysis of MERS-CoV transmission pointing to repeated spillover from camels to humans (*11*). Furthermore, these data demonstrate that the mutations observed in MERS-CoV strains sequenced to date do not markedly dampen plasma neutralizing activity.

To assess neutralization breadth against a phylogenetically distant merbecovirus, we evaluated inhibition of a VSV pseudotyped with the S glycoprotein of the recently described HKU4-related pangolin isolate (MjHKU4r-CoV-1). MjHKU4r-CoV-1 S shares 65.4% amino acid sequence identity with MERS-CoV EMC/2012 S, including strict conservation of 8 out of 16 receptor-binding motif residues, likely explaining retention of human DPP4 utilization (*61*). Only 1 of 30 plasma samples tested had detectable but weak neutralization potency against MjHKU4r-CoV-1 S VSV (**Figure 1E**; **Figure S4**). These results suggest that although a single MERS-CoV infection elicits neutralizing antibodies against known circulating MERS-CoV variants, it is insufficient to induce potent pan-merbecovirus neutralizing activity.

### MERS-CoV S_1_-directed antibodies account for most plasma neutralizing activity

Given that S_1_-directed plasma antibodies, particularly those targeting the RBD, account for virtually all SARS-CoV-2 neutralizing activity upon infection or vaccination (*44, 46*), we sought to determine if MERS-CoV infection-elicited antibodies share a similar specificity. IgG binding titers against the MERS-CoV EMC/2012 S_1_ subunit ranged from ≤10 to 20,101 (GMT: 963; **Figure 2A**; **Figure S5**) for the 29 plasma samples evaluated and positively correlated with neutralizing activity (Spearman r = 0.65; **Figure 2B**), as observed with S-directed IgG binding titers.

**Figure 2.**
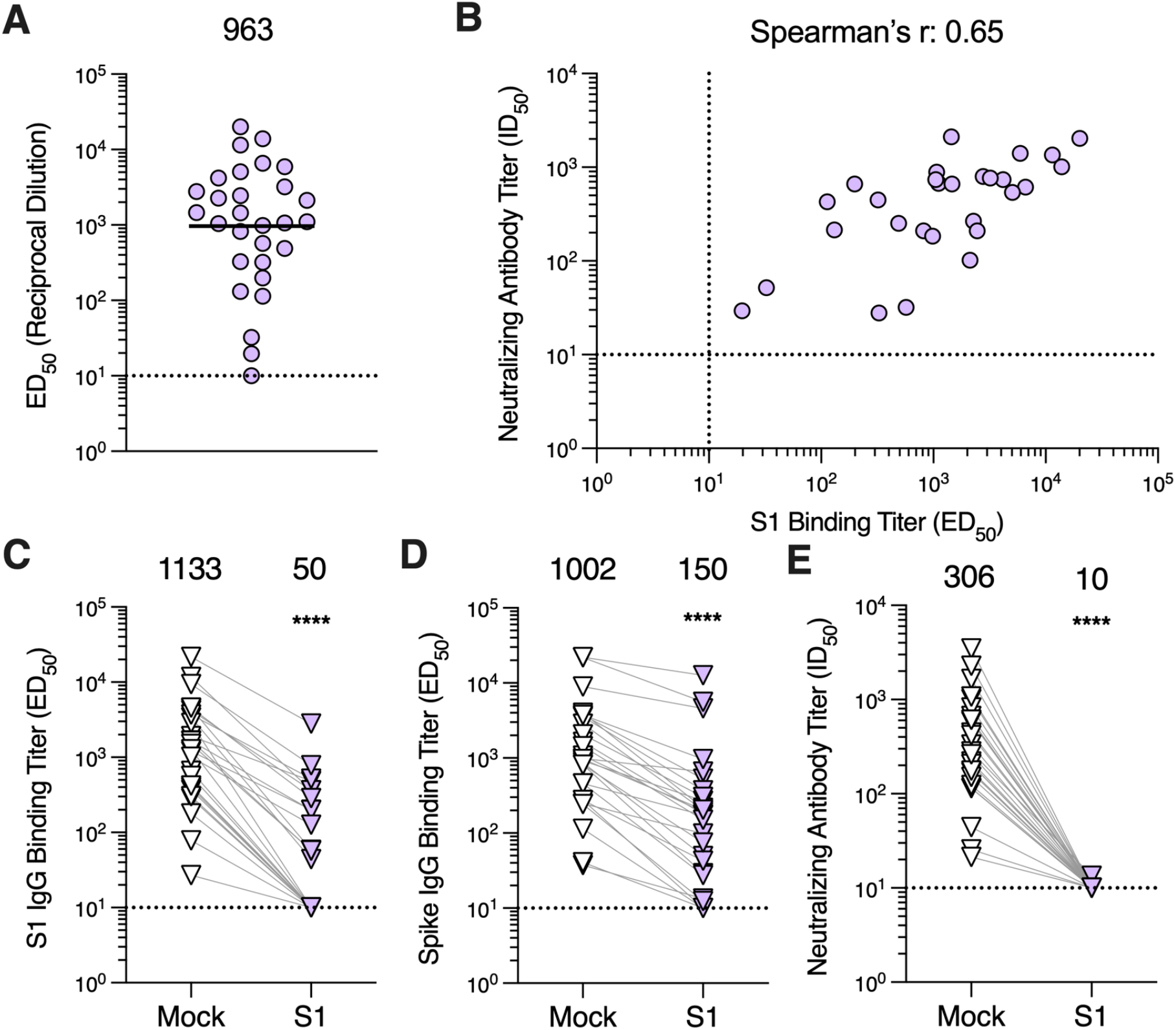
Antibodies directed against the S_1_ subunit of the MERS-CoV spike protein are responsible for nearly all neutralizing activity of plasma. **A)** IgG binding titers (ED_50_) against the MERS-CoV EMC/2012 S_1_ subunit measured for plasma samples. The sample with the highest S binding titer per individual was included in the analysis with 29 samples being included in total. The GMT is represented by the black bar and displayed above the plot. The limit of detection (ED_50_: 10) of the assay is represented by the dashed line. Data reflect results obtained from one biological replicate and are representative of data obtained from at least two biological replicates conducted with unique batches S_1_ protein. **B)** Correlation analysis between S_1_ IgG binding titers and neutralizing antibody titers for samples with detectable S_1_ binding titers. **C)** S_1_ and **D)** S IgG binding titers and **E)** neutralizing antibody titers in mock-depleted and S_1_-depleted plasma samples. One sample per individual was included in the analysis with 27 samples being included in total. Mock- and S_1_-depleted samples from the same individual are connected with a gray line. GMTs are displayed above the plot and the limit of detection (ED_50_ or ID_50_: 10) is represented by the dashed line. Data reflect results obtained from one biological replicate and are representative of data obtained from at least two biological replicates conducted with unique batches of S_1_ and S proteins and pseudovirus. Comparisons between mock-and S1-depleted groups were made using the Wilcoxon matched-pairs signed rank test. ****p < 0.0001.

To further understand the contribution of S_1_-directed antibodies to plasma neutralizing activity, we depleted the plasma samples of S_1_-directed antibodies by incubating plasma with magnetic beads coupled to the MERS-CoV S_1_ subunit. Following incubation with the S_1_-coated beads, S_1_ IgG binding titers were undetectable for 13 out of 27 samples (ED_50_ ≤10) and reduced 74.5 to 98.7% for the remaining 14 samples, confirming successful depletion (**Figure 2C**; **Figure S6A**). Furthermore, we observed a 89.5% reduction in S-directed IgG binding titers, with 3 out of 27 samples reaching the limit of detection (ED_50_ ≤ 10), corresponding to a statistically significant difference in GMTs of 150 and 1,002 for S_1_-depleted and mock-depleted plasma samples, respectively (p = 0.0002; **Figure 2D**; **Figure S6B**). S_1_-directed antibodies therefore account for the majority of S binding antibodies in plasma, underscoring the greater immunogenicity of the S_1_ subunit compared to the S_2_ subunit. Furthermore, depletion of S_1_-directed antibodies resulted in a near complete loss of neutralizing activity with only a single sample retaining detectable neutralization (ID_50_ > 10; **Figure 2E**; **Figure S6C**), which was, however, reduced by 95.2%. These data show that S_1_-directed antibodies account for nearly all polyclonal plasma neutralizing activity in plasma samples post MERS-CoV infection.

### MERS-CoV RBD-directed antibodies account for most plasma neutralizing activity

As the S_1_ subunit comprises at least two domains targeted by neutralizing antibodies, namely the RBD and the NTD (*14*), we set out to understand the contribution of antibodies directed against each of these domains to polyclonal plasma neutralizing activity. RBD-directed IgG binding titers ranged from ≤10 to 14,253 (GMT: 1,007; **Figure 3A**; **Figure S7A**) and NTD-directed IgG binding titers ranged from ≤10 to 821 (GMT: 45; **Figure 3B**; **Figure S7B**) and both of them positively correlated with neutralizing antibodies titers (Spearman r = 0.76 and 0.50, respectively; **Figure 3C**), concurring with the fact that the S_1_ subunit is the main target of neutralizing antibodies.

**Figure 3.**
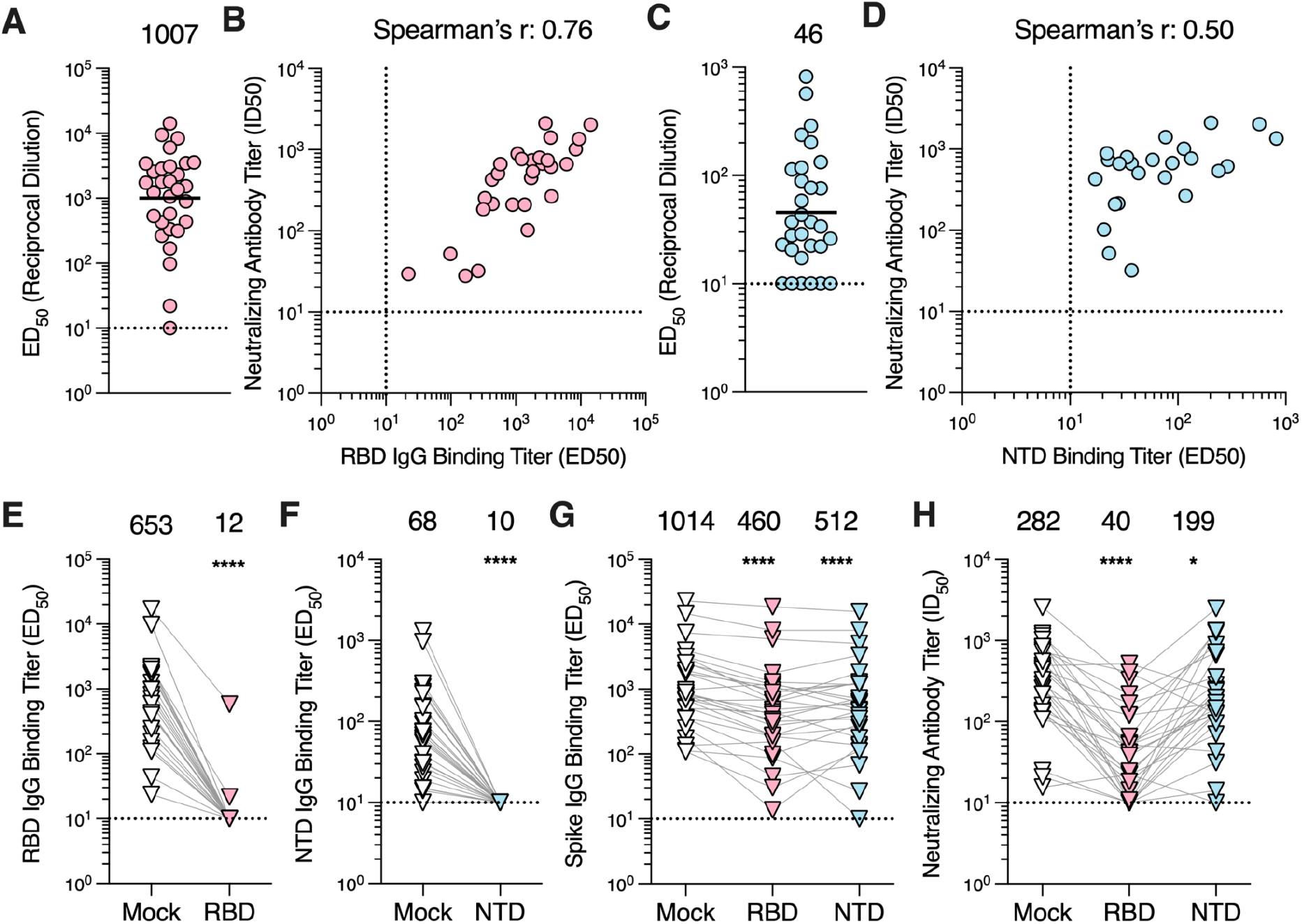
RBD-directed antibodies account for the majority of neutralizing activity in plasma. **A)** IgG binding titers (ED_50_) against the MERS-CoV EMC/2012 RBD measured for the plasma samples. **B)** Correlation analysis between RBD IgG binding titers and neutralization potency of the plasma samples with detectable RBD binding titers. **C)** IgG binding titers (ED_50_) against the MERS-CoV EMC/2012 NTD determined for the plasma samples. **D)** Correlation analysis between NTD IgG binding titers and neutralizing antibody titers in the plasma samples with detectable NTD binding titers. For the RBD and NTD ELISAs, the sample with the highest S binding titer per individual was included in the analysis. The GMT is represented by the black bar and presented above the plot. The limit of detection (ED_50_: 10) of the assay is indicated by the dashed line. Data reflect results obtained from one biological replicate and are representative of data obtained from at least two biological replicates conducted with unique batches of RBD or NTD protein. **E)** RBD IgG binding titers for mock- and RBD-depleted plasma samples. **F)** NTD IgG binding titers for mock- and NTD-depleted plasma samples. **G)** S IgG binding titers and **H)** neutralizing antibody titers for mock-, RBD-, and NTD-depleted plasma. Six of the 28 RBD-depleted plasma and 1 of the 28 NTD-depleted plasma samples exhibited neutralizing titers below the limit of detection. The one sample with the highest S binding titer per individual was included in the analysis. Data presented are from one biological replicate and reflective of two biological replicates completed with unique batches of RBD, NTD, and S proteins as well as distinct batches of pseudovirus. GMTs are displayed above the plots and the limit of detection (ED_50_ or ID_50_: 10) is indicated by the dashed line. Comparisons between RBD or NTD IgG binding titers of the mock- and RBD or NTD-depleted groups, respectively, were made using the Wilcoxon matched-pairs signed. Comparison between the spike IgG bindings titers and neutralizing antibody titers of mock-, RBD- and NTD-depleted groups were made using Dunn’s multiple comparison test. *p<0.05; ****p < 0.0001.

To further resolve the contribution of RBD-directed and NTD-directed antibodies to neutralizing activity, we depleted the plasma samples of domain-specific antibodies using RBD- or NTD-coated magnetic beads. Following depletion, 26 out of 28 plasma samples had RBD-directed IgG binding titers below the limit of detection (ED_50_ ≤ 10; **Figure 3E**), with 96.6 and 99.0% reductions for the remaining 2 samples, whereas none of the plasma samples had detectable NTD-directed IgGs (**Figure 3F**). Furthermore, we observed a significant reduction in S-directed IgG binding titers for plasma samples depleted of RBD-directed antibodies (54.6%, p≤ 0.0001; GMT: 460) and of NTD-directed antibodies (49.5%, p ≤ 0.0001; GMT: 512) compared to mock-depleted plasma (GMT: 1,014, **Figure 3G**). The residual neutralization potency of these plasma samples was significantly reduced after depletion of RBD-directed antibodies (85.8%, GMT: 40, p = <0.0001) or of NTD-directed antibodies (29.4%, GMT: 199, p = 0.0151) compared to the mock-depleted samples (GMT: 282, **Figure 3H**). Collectively, these data suggest that both the RBD and the NTD are targeted by neutralizing antibodies elicited by MERS-CoV infection with RBD-directed antibodies accounting for most of the polyclonal plasma neutralizing activity.

### Fusion machinery-directed neutralizing antibodies are rare post MERS-CoV infection

To determine the antigenic sites targeted by infection-elicited polyclonal plasma antibodies, we carried out competition ELISAs using a panel of mAbs recognizing distinct MERS-CoV S epitopes in the RBD, NTD, and S_2_ subunit (fusion machinery). We examined the magnitude of competition of binding to MERS-CoV S between plasma and the RBD-directed mAbs LCA60, S41, and 4C2, along with the human DPP4 receptor (**Figure 4A**). LCA60 binds the saddle of the RBD, overlapping partially with the DPP4 binding site (*29, 30*). The S41 epitope overlaps entirely with the DPP4-binding site although it has a smaller footprint (*31*). 4C2 binds an RBD epitope partially overlapping with the RBM but that remains exposed in the closed S trimer, (*34*). Out of the 30 plasma samples tested, 37% (n = 11) competed with DPP4 binding (half-maximum blocking titers, BD50 > 10) whereas 33% (n = 10), 20% (n = 6) and 53% (n = 16) competed with the LCA60, S41 and 4C2 mAbs, respectively (**Figure 4B**). Antibody responses elicited by MERS-CoV infection thus more prevalently target RBD antigenic sites that remain fully exposed in the closed S trimer relative to epitopes that are fully or partially occluded.

**Figure 4.**
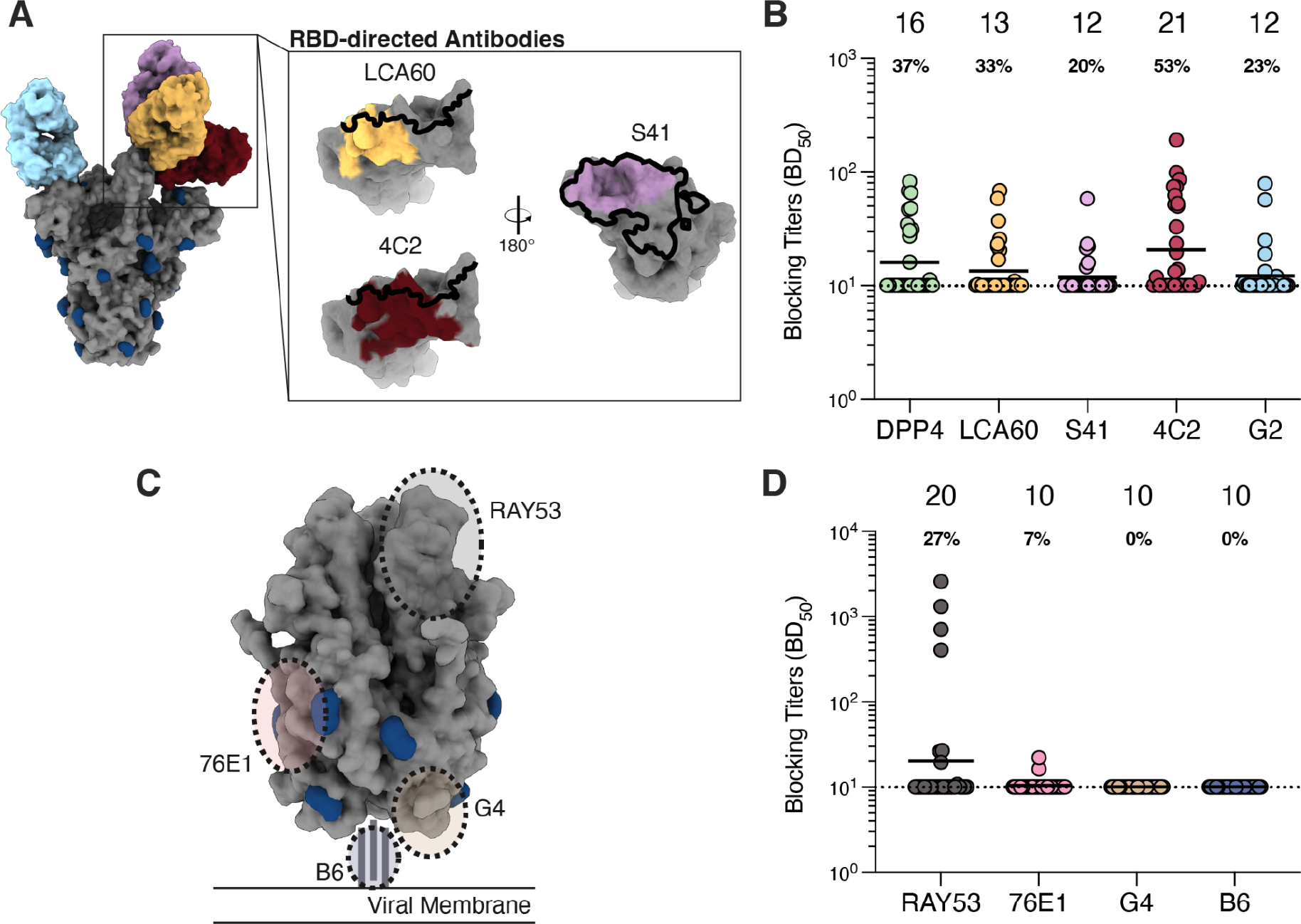
Plasma antibodies target S_1_ neutralizing epitopes, but rarely S_2_ epitopes. **A)** Epitopes targeted by S_1_-directed antibodies included for the competition ELISAs. The RBM is outlined in black and glycans are represented in dark blue. **B)** Blocking titers (half-maximal blocking dilution [BD_50_]) measured for plasma samples against the indicated S_1_-directed antibodies. **C)** Epitopes targeted by S_2_-directed antibodies included in the competition ELISAs. Glycans are displayed in dark blue. **D)** Blocking titers (BD_50_) in plasma samples against the specified S_2_-directed antibodies. The sample with the highest S binding titer per individual was analyzed. GMTs are displayed above the plot (indicated by black bar) and the portion of individuals exhibiting detectable blocking titers is displayed below the GMT for each antibody. Data presented are from one biological replicate and representative of data collected from at least two biological replicates conducted with distinct batches of antibodies and biotinylated S protein.

We further examined the capacity of plasma to block binding to MERS-CoV S of the NTD-directed neutralizing antibody G2, which binds to an epitope on the viral membrane distal side of the NTD (**Figure 4A**) (*43*). Only 23% (n = 7) of the plasma samples competed with G2 despite the retained exposure of this antigenic site irrespective of S conformation (**Figure 4B**). The low frequency of G2-blocking antibodies in plasma is consistent with the low NTD-directed binding titers (**Figure 3C**) and with the depletion data, which suggest that NTD-directed antibodies make a smaller overall contribution to plasma neutralizing activity, relative to RBD-directed antibodies.

We next quantified competition of plasma against a panel of S_2_-directed mAbs including RAY53, 76E1, G4, and B6 (Figure 4C). RAY53 is a weakly neutralizing mAb that binds to the hinge region of the S_2_ subunit in the prefusion conformation (*39*). 76E1 targets the fusion peptide and broadly reacts with both alphacoronaviruses and betacoronaviruses (*38*). G4 binds to a variable loop in the MERS-CoV S_2_ connector domain (*36*). B6 targets the S stem helix and cross-reacts with all human-infecting betacoronaviruses (albeit weakly with HKU1) (*40*). 27% (n = 8) of samples competed with RAY53 binding (BD_50_ > 10; Figure 4D) and 4 of them had RAY53-blocking titers greater than those observed for the RBD- and NTD-directed antibodies (BD_50_ values ranging from 404 to 2,588). We speculate that RAY53 binding may be hindered by some RBD-directed antibodies, particularly those that lock S in the closed conformation, as observed for some SARS-CoV-2 mAbs (*62*). For the remaining three S_2_-directed mAbs, 7% (n= 2) of plasma samples had detectable competition with 76E1 and none of the 30 samples tested had detectable competition with G4 or B6. These findings coupled with the S_1_-antibody depletion data suggest that MERS-CoV infection does not induce robust binding and neutralizing antibody responses against the MERS-CoV S_2_ subunit (Figure 2 D-E).

## Discussion

Here, we examined antibody responses in plasma samples obtained from individuals hospitalized with MERS-CoV infections between 2017-2019. We observed S binding titers and neutralizing antibody titers peaking between 1 to 6 weeks post symptom onset or hospitalization. We found that plasma antibodies neutralize MERS-CoV variants with similar potencies, but display minimal neutralizing activity against the highly divergent merbecovirus MjHKU4r-CoV-1. We further showed that the S_1_ subunit is immunodominant with S_1_-directed antibodies accounting for the majority of the neutralization potency of plasma. Moreover, we observed that both the RBD and NTD contribute to the S-directed IgG binding activity of plasma, but RBD-directed antibodies were responsible for most of the neutralizing activity. Finally, we used a panel of structurally characterized neutralizing mAbs to show that plasma antibodies target neutralizing epitopes on the S_1_ subunit, but rarely those on the S_2_ subunit.

MERS-CoV infection induced heterogeneous humoral immune responses among individuals in terms of magnitude and durability of S-directed IgG binding and neutralizing antibodies. These data are consistent with previous observations on the immune response developed after SARS-CoV-2 infection (*44, 46, 63*). As underlying comorbidities likely influenced the magnitude and durability of the antibody responses, vaccination will likely be required to generate a homogeneously robust and durable response in populations at risk of MERS-CoV infection, as observed upon SARS-CoV-2 vaccination (*46*). MERS-CoV variants identified to date have accumulated relatively few S mutations to date similar to the earliest SARS-CoV-2 variants, including CAL.20C (B.1.427/B.1429) and Delta (B.1.617.2) (*64, 65*). Consistently, the degree of antibody evasion by the MERS-CoV variants is similar to that of these SARS-CoV-2 variants that emerged early in the COVID-19 pandemic.

Prior studies of plasma antibody responses elicited against SARS-CoV-2 S following either SARS-CoV-2 infection or vaccination with S-encoding mRNA vaccines demonstrated that the RBD is immunodominant and accounts for the majority of plasma neutralizing activity (*44, 46–49*). These studies further indicated that NTD-directed antibodies contribute to plasma neutralizing activity against infection- or vaccine-matched variants, albeit to a lesser extent than RBD-directed antibodies (*46, 66*). Our results on the contribution of MERS-CoV S domains to the neutralizing activity of plasma are consistent with observations made with SARS-CoV-2 infection- and vaccine-elicited plasma samples. As RBD-directed antibodies are responsible for the majority of plasma neutralizing activity for both SARS-CoV-2 and MERS-CoV, we speculate RBD-directed antibodies will account for most of the plasma neutralization activity for all betacoronaviruses. As such, betacoronavirus RBD-based vaccines will likely confer high neutralizing antibody titers and robust protection across this genera and could become an universal strategy to develop betacoronavirus vaccines, as successfully implemented with the SKYcovione COVID-19 vaccine (*67, 68*).

Our plasma epitope mapping analysis indicated that the ridge of the RBM exposed in the closed S trimer is frequently targeted by plasma antibodies, while RBD epitopes partially or fully occluded in the closed S conformation are less frequently targeted. These data for MERS-CoV plasma are consistent with epitope mapping data for SARS-CoV-2 plasma, which suggested the analogous site on the SARS-CoV-2 RBD (site Ib) is frequently targeted by plasma antibodies whereas occluded antigenic sites (sites IIa, IIb, and IIc) are less commonly targeted (*44*). However, as far fewer neutralizing RBD-directed antibodies for MERS-CoV (*29–32, 34*) have been characterized than for SARS-CoV-2 (*44, 69, 70*), it is likely that there are additional neutralizing epitopes on the MERS-CoV RBD that contribute to the neutralizing activity of plasma. Additional studies focused on discovering and characterizing antibodies targeting the MERS-CoV RBD will further resolve immunodominant epitopes as well as elucidate those epitopes that are resilient to viral evolution and contribute to the cross-neutralization of related merbecoviruses (*42*).

We observed minimal contribution of S_2_ subunit-directed antibodies to polyclonal plasma neutralization as known epitopes targeted by neutralizing mAbs are rarely targeted by plasma antibodies, concurring with the rarity of stem helix- and fusion peptide-directed antibodies upon SARS-CoV-2 infection or vaccination (*41, 71*). We speculate that the lack of plasma antibodies targeting S_2_ epitopes may be due to inaccessibility until the S_1_ subunit binds DPP4 or is shed, as is the case for 76E1 and RAY53 (*38, 39*), or being located close to the viral membrane and likely less accessible to recognition by B cells, as is the case for G4 and B6 (*36, 40*). While the development of S_2_ subunit antigens has been a recent goal for SARS-CoV-2 vaccine design due to the resiliency of S_2_-directed antibodies to the rapid evolution of SARS-CoV-2 (*72–78*), similar vaccine antigens may not be necessary for MERS-CoV. Our demonstration of the retention of plasma neutralizing activity against human and camel MERS-CoV strains along with the relatively few S_1_ mutations in MERS-CoV strains sequenced to date (*11, 60*) suggest vaccination with prefusion S or RBD-based vaccines may be sufficient for resiliency against MERS-CoV evolution, concurring with data presented in our companion manuscript.

In conclusion, our data demonstrate MERS-CoV RBD-directed antibodies account for the majority of neutralizing activity of plasma, consistent with similar studies on SARS-CoV-2 plasma, and suggest vaccines in development for MERS-CoV and other betacoronaviruses should focus on eliciting a robust RBD-directed neutralizing antibody response.

## Acknowledgements

This study was supported by the National Institute of Allergy and Infectious Diseases (P01AI167966, DP1AI158186 and 75N93022C00036 to D.V.), an Investigators in the Pathogenesis of Infectious Disease Awards from the Burroughs Wellcome Fund (D.V.). D.V. is an Investigator of the Howard Hughes Medical Institute and the Hans Neurath Endowed Chair in Biochemistry at the University of Washington.

## Author Contributions

AA and DV conceived the study and designed the experiments; AA and CS recombinantly expressed and purified glycoproteins. AA performed all other experiments. AYA, MAM, ZAM and AbA collected the MERS-CoV clinical samples and prepared plasma. AA and D.V. analyzed the data and wrote the manuscript with input from all authors. DV supervised the project.

## Methods

### Cell culture

Expi293 cells were grown in Expi293 media at 37°C and 8% CO_2_ rotating at 130 RPM. HEK-293T cells were grown in DMEM supplemented with 10% FBS and 1% PenStrep at 37°C at 5% CO_2_. Vero E6 cells stably expressing the human protease TMPRSS2 (Vero-TMPRSS2) were grown in DMEM supplemented with 10% FBS, 1% PenStrep and 8 µg/mL puromycin at 37°C and 5% CO_2_ (*79*).

### Plasma donors and selection of plasma samples

Samples were collected from patients with confirmed diagnosis of MERS admitted to the critical care unit in Prince Mohammed Bin Abdulaziz Hospital in Riyadh, Saudi Arabia. Family members responsible for patients who agreed to participate in the study provided informed consent for serial blood collections from patients. The study was approved by the institutional research board of King Faisal specialist hospital and research center in Jeddah, Saudi Arabia IRB 2017-73. Demographic data for the individuals is presented in Table S1.

For neutralization assays with variant MERS-CoV and MjHKU4r-CoV-1 pseudoviruses, RBD, NTD, and S_1_ ELISAs and depletion experiments, and mAb competition ELISAs, one sample per individual was included in each analysis. We included the sample with the highest S binding titer for each individual, except for individual 27 where the sample with the second highest S binding titer was included due to volume constraints. For the depletion experiments, we excluded samples from individuals 2 and 31 due to low binding and neutralizing antibody titers. For both the S_1_ ELISA and depletion experiments, individual 22 was excluded due to plasma volume limitations. The same set of plasma samples were included across the aforementioned assays.

### Constructs

The construct encoding the prefusion stabilized MERS-CoV spike (S2P) ectodomain was previously described (*18, 30*). Constructs encoding the B6 heavy and light chains were previously described (*40*), those encoding the G2 and G4 heavy and light chains were gifted by Jason McLellan (*36, 43*), those encoding the RAY53 heavy and light chains were gifted by Jennifer Maynard (*39*), and the construct encoding the full-length MERS-CoV EMC/2012 spike protein was gifted by Gary Whittaker (*13*).

The full-length MERS-CoV United Kingdom/H123990006/2012, MERS-CoV Hu/Riyadh-KSA-3181/2015, MERS-CoV Camel/Kenya/M23C14/2019, MERS-CoV Korea/Seoul/168-1-2015 and MjHKU4r-CoV-1 Δ16 spike proteins were codon optimized, synthesized, and inserted into pcDNA3.1(+) by Genscript. The heavy chain Fab sequences for LCA60, S41, 4C2, and 76E1 fused to an N-terminal µ-phosphatase or mouse Ig heavy signal peptide sequence and C-terminal human FC tag were codon optimized, synthesized, and inserted into pcDNA3.1(+) by Genscript. The light chain Fab sequence for these antibodies with an N-terminal µ-phosphatase or mouse Ig heavy signal peptide sequence were codon optimized, synthesized, and inserted into pcDNA3.1(+) by Genscript. The MERS-CoV NTD (1–357) with a C-terminal octa-his tag, MERS-CoV RBD (382–588) with an N-terminal µ-phosphatase signal peptide and C-terminal thrombin cleavage site followed by an octa-his tag, and MERS-CoV S1 (1–747) with a C-terminal octa-his tag were codon optimized, synthesized, and inserted into pcDNA3.1(+), pCMVR, and pcDNA3.4, respectively, by Genscript. The human DPP4 ectodomain (39–766) with a N-terminal CD5 leader sequence and C-terminal human FC tag was synthesized and inserted into pcDNA3.1(-).

### Recombinant protein expression and purification

The MERS-CoV RBD, MERS-CoV NTD, and MERS-CoV S_1_ were expressed and purified as previously described (*18, 30, 80–82*). In brief, Expi293 cells were grown to a density of 3 x 10^6^ cells/mL and transfected using the Expifectamine293 transfection kit following the manufacturer’s recommendations. Four to five days following transfection, the supernatants were collected, clarified by centrifugation, and flowed over HisTrap FF or HP affinity columns. The columns were then washed with ten column volumes of 20 mM imidazole, 25 mM sodium phosphate pH 8.0, and 300 mM NaCl after which the proteins were eluted using a gradient up to 500 mM imidazole. The proteins were then buffer exchanged into 20 mM sodium phosphate pH 8.0 and 100 mM NaCl and concentrated using centrifugal filters, flash frozen, and stored at -80°C until use.

The MERS-CoV S2P was expressed and purified similar to above. Following elution from the affinity column, the protein was further purified by size-exclusion chromatography using a Superose 6 Increase 10/300 GL column. The protein was concentrated, flash frozen, and stored at -80°C. For the biotinylated MERS-CoV S2P, MERS-CoV S2P was expressed and purified as described above. After elution from the affinity column, the protein was buffer exchanged into 25 mM Tris-HCl pH 8.0 and 150 mM NaCl. The purified spike protein was biotinylated using the BirA biotin-protein ligase reaction kit (Avidity) following the manufacturer’s recommendation. The biotinylated protein was re-purified using an HisTrap HP column followed by size-exclusion chromatography using a Superose 6 Increase 10/300 GL column. The protein was concentrated, flash frozen, and stored at -80°C.

Recombinant monoclonal antibodies were produced by transfecting Expi293 cells at a density of 3 x 10^6^ cells/mL with equal masses of the heavy and light chain constructs using Expifectamine293. Four to five days following transfection, the supernatants were harvested and clarified by centrifugation. The recombinant antibodies were captured using HiTrap Protein A HP affinity columns after which the columns were washed with ten column volumes of 20 mM sodium phosphate pH 8.0 and the proteins were eluted 0.1 M citric acid pH 3.0, which was neutralized with 1 M Tris-HCl pH 9.0. The antibodies were buffer exchanged into 20 mM sodium phosphate pH 8.0 and 100 mM NaCl and concentrated using centrifugal filters and stored at 4°C until use. The human DPP4 ectodomain was expressed and purified as described above, flash frozen, and stored at -80°C until use.

### Enzyme-linked immunosorbent assay (ELISA)

The MERS-CoV S2P, RBD, NTD, or S_1_ recombinant proteins were diluted to 0.003 mg/mL and added to 384-well Maxisorp plates overnight at room temperature. The following day, plates were slapped dry and blocked with Blocker Casein for 1 hour at 37°C. The plates were slapped dry and plasma samples diluted in Tris-buffered saline with 0.1% Tween 20 (TBST) at a starting dilution of 1:10 to 1:270 and serially diluted 1:3 thereafter were added to the plates. The plates were incubated for 1 hour at 37°C, slapped dry, and washed four times with TBST. A goat anti-human IgG (H+L) HRP conjugated antibody (diluted 1:5,000 in TBST; Seracare) was added to each well. The plates were again incubated for 1 hour at 37°C, slapped dry, and washed four times with TBST. SureBlue Reserve TMB 1-Component Microwell Peroxidase Substrate was added to each well and allowed to develop for 90 seconds after which an equal volume of 1 N HCl was added to quench the reaction. The absorbance at 450 nm was immediately measured using a BioTek Synergy Neo2 plate reader. The resulting data were analyzed in GraphPad Prism 10 using a four parameter logistic curve to determine the ED_50_ for each sample. At least two biological replicates using two distinct batches of recombinant protein were performed for each sample and antigen.

### Pseudotyped VSV production

VSV pseudotyped with the MERS-CoV or MjHKU4r-CoV-1 spike proteins were produced as previously described (*80–83*). In brief, HEK293T cells were seeded onto poly-D-lysine coated 10 cm^2^ plates at a density of 5 x 10^6^ cells and grown overnight until they reached approximately 80-90% confluency. The following day, the growth media was exchanged to DMEM containing 10% FBS and the cells were transfected with the spike constructs using Lipofectamine 2000. After 20 to 24 hours, the cells were washed three times with DMEM and infected with VSVΔG/luc. Two hours after infection, the cells were washed five times with DMEM and left overnight in DMEM supplemented with an anti-VSV-G antibody (I1-mouse hybridoma supernatant diluted 1:25, from CRL-2700, ATCC). Twenty to 24 hours later, the supernatant was harvested, clarified by centrifugation at 4,200 RPM for 10 minutes, filtered using a 0.45 µm filter, and concentrated with a 100 kDa filter (Amicon). The resulting pseudovirus was frozen at -80°C until use.

### Pseudovirus neutralization assays

Neutralization assays were performed as previously described (*80–83*). In brief, Vero-TMPRSS2 cells were split in white-walled, clear bottom plates at a density of 18,000 cells per well. The cells were grown overnight until they reached 90-95% confluency. Plasma samples were diluted in DMEM starting at a 1:10 dilution and serially diluted 1:3 thereafter and mixed with an equal volume of pseudotyped VSV diluted 1:50 to 1:100 in DMEM. The virus-plasma mixtures were incubated for 30 minutes at room temperature after which the growth media was removed from the Vero-TMPRSS2 cells and replaced with the virus-plasma mixture. The cells were incubated for 2 hours at 37°C after which an equal volume of DMEM supplemented with 20% FBS and the 2% PenStrep was added to each well. Twenty to 24 hours later, ONE-Glo EX was added directly to each well, the plates were then incubated for 5 minutes at 37°C, and the luminescence values were recorded using a BioTek Synergy Neo2 plate reader. The data were normalized in GraphPad Prism 10 using the relative light unit (RLU) values measured for uninfected cells to define 0% infectivity and RLU values recorded for cells infected with pseudovirus without plasma to define 100% infectivity. ID_50_ values for plasma samples were determined from the normalized data using a [inhibitor] vs. normalized response – variable slope model using two technical replicates to generate the curve fits. At least two biological replicates using distinct batches pseudoviruses were performed for each sample.

### Depletion Assays

Dynabeads His-Tag Isolation and Pulldown beads (ThermoFisher) were washed once with Tris-buffered saline (TBS) containing 0.01% Tween 20. His-tagged MERS-CoV RBD, NTD, or S_1_ was added to the magnetic beads at a 1.5-fold mass excess relative to the reported binding capacity of the beads. The beads and proteins were incubated for 30 minutes at room temperature with constant rotation. Unbound protein was then removed by washing the beads three times with TBS with 0.01% Tween 20. The antigen coated beads were resuspended in TBS with 0.01% Tween 20 and stored at 4°C until use.

Plasma samples were mixed with the RBD, NTD or S_1_ coated beads in TBS with 0.01% Tween 20 at a 1:4 volume ratio and incubated at 37°C for 1 hour. Following this incubation, the supernatant was transferred to a new tube that contained an additional 4 volumes of beads. The mixture was incubated for an additional hour at 37°C after which the supernatant was transferred to a new tube. A mock depletion was performed concurrently following the same protocol as above except the plasma was incubated with uncoated beads. ELISAs against the depleting antigen and MERS-CoV spike and neutralization assays against VSV pseudotyped with the MERS-CoV EMC/2012 spike were performed as described above using the depleted plasma as input. The starting 1:10 dilution factor incorporates the dilutions performed to deplete the plasma of the domain-specific antibodies. At least two biological replicates using distinct batches of proteins and pseudoviruses were performed for each sample.

### Competition ELISA

To determine the EC_50_ of the antibodies used in the competition ELISAs, the antibodies were diluted to 0.003 mg/mL, plated on 384-well Maxisorp plates, and incubated overnight at room temperature. The following day, the plates were slapped dry and blocked with Blocker Casein at 37°C for 1 hour. The plates were again slapped dry and biotinylated MERS-CoV S2P diluted to 0.3 mg/mL and serially diluted 1:3 in TBST thereafter was added to each well. The plates were incubated for 37°C for 1 hour, slapped dry, and washed four times with TBST. Ultra streptavidin-HRP diluted 1:2,000 in TBST was added to each well after which the plates were again incubated at 37°C for 1 hour, slapped dry, and washed four times with TBST. SureBlue Reserve TMB 1-Component Microwell Peroxidase Substrate was added to each well and allowed to develop for 120 seconds after which an equal volume of 1 N HCl was added to quench the reaction. The absorbance at 450 nm was immediately measured using a BioTek Synergy Neo2 plate reader. The resulting data were analyzed in GraphPad Prism 10 using a four parameter logistic curve to determine the EC_50_ for each antibody.

For the competition ELISAs, 384-well Maxisorp plates were coated with the monoclonal antibodies and blocked as described above. Plasma samples were diluted in non-binding 384-well plates beginning with a 1:5 dilution in TBST followed by a 1:3 serial dilution thereafter. Biotinylated MERS-CoV S2P was diluted to a concentration twice the determined EC_50_ in TBST and added to each well of the non-binding plates. The plasma-spike mixtures were incubated for 1 hour at 37°C after which the plasma-spike mixtures were transferred to monoclonal antibody coated plates. The plates were incubated for 1 hour at 37°C, slapped dry, and washed four times with TBST. Ultra streptavidin-HRP diluted 1:2,000 in TBST was added to the plates and the plates were incubated for 1 hour at 37°C, slapped dry, and washed four times with TBST. SureBlue Reserve TMB 1-Component Microwell Peroxidase Substrate was added to each well and allowed to develop for 120 seconds after which an equal volume of 1 N HCl was added to quench the reaction. The absorbance at 450 nm was immediately measured using a BioTek Synergy Neo2 plate reader. The resulting data was normalized using the absorbance values from wells without spike added to define 0% binding and the absorbance values from wells with spike but no plasma added to define 100% binding. BD_50_ values were determined using the [inhibitor] vs. normalized response – variable slope model using two technical replicates to generate the curve fits. Two biological replicates using distinct batches of spike protein and monoclonal antibodies were performed for each sample.

## Supplemental Figures

**Figure S1.**
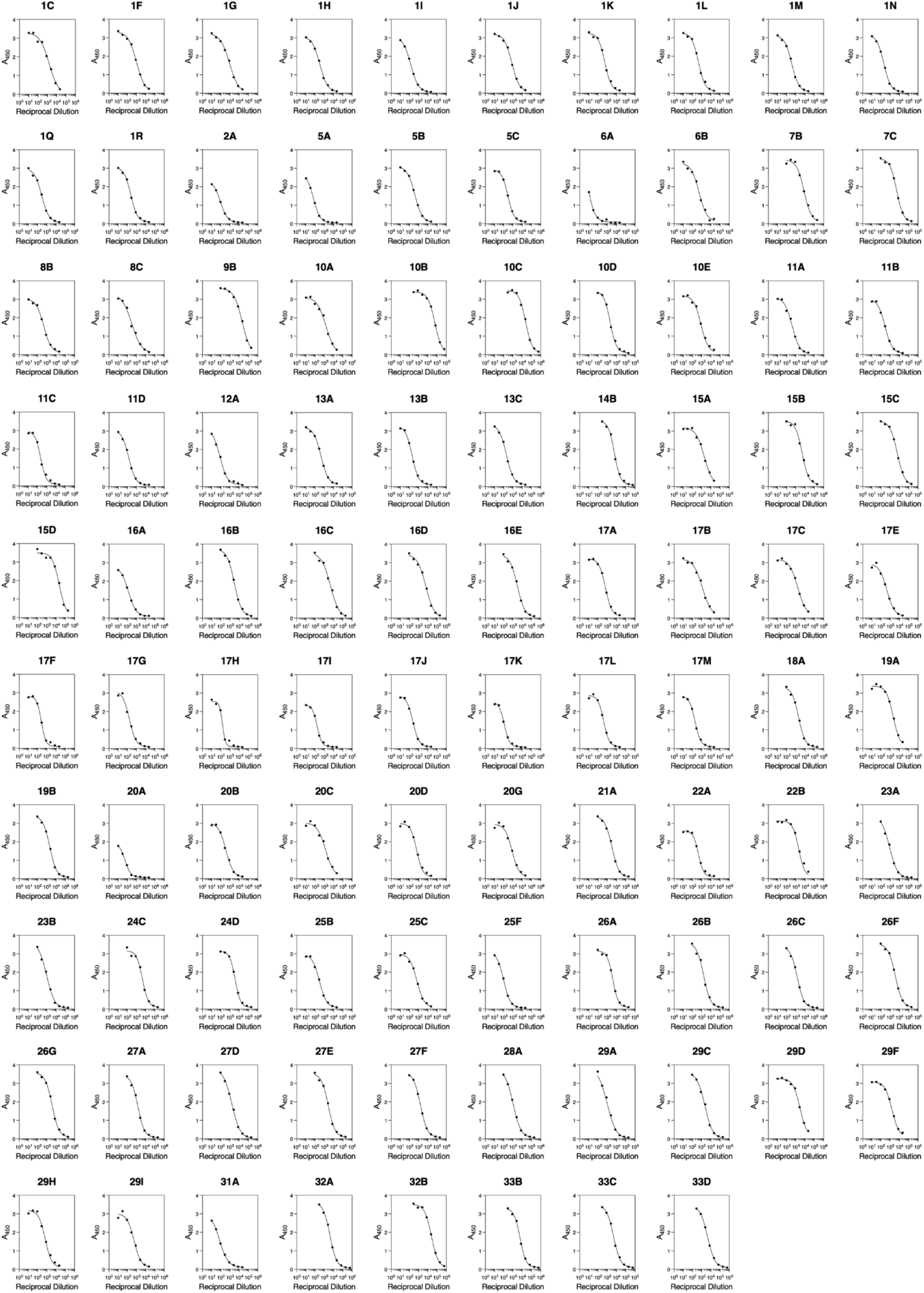
Evaluation of MERS-CoV S-directed plasma IgG binding titers. Dose-response curves of plasma IgG binding to prefusion-stabilized MERS-CoV EMC/2012 2P S for each of the 98 samples analyzed in this study by ELISA. Data are presented for one representative biological replicate. At least two biological replicates each using a unique batch of MERS-CoV S were conducted for each sample.

**Figure S2.**
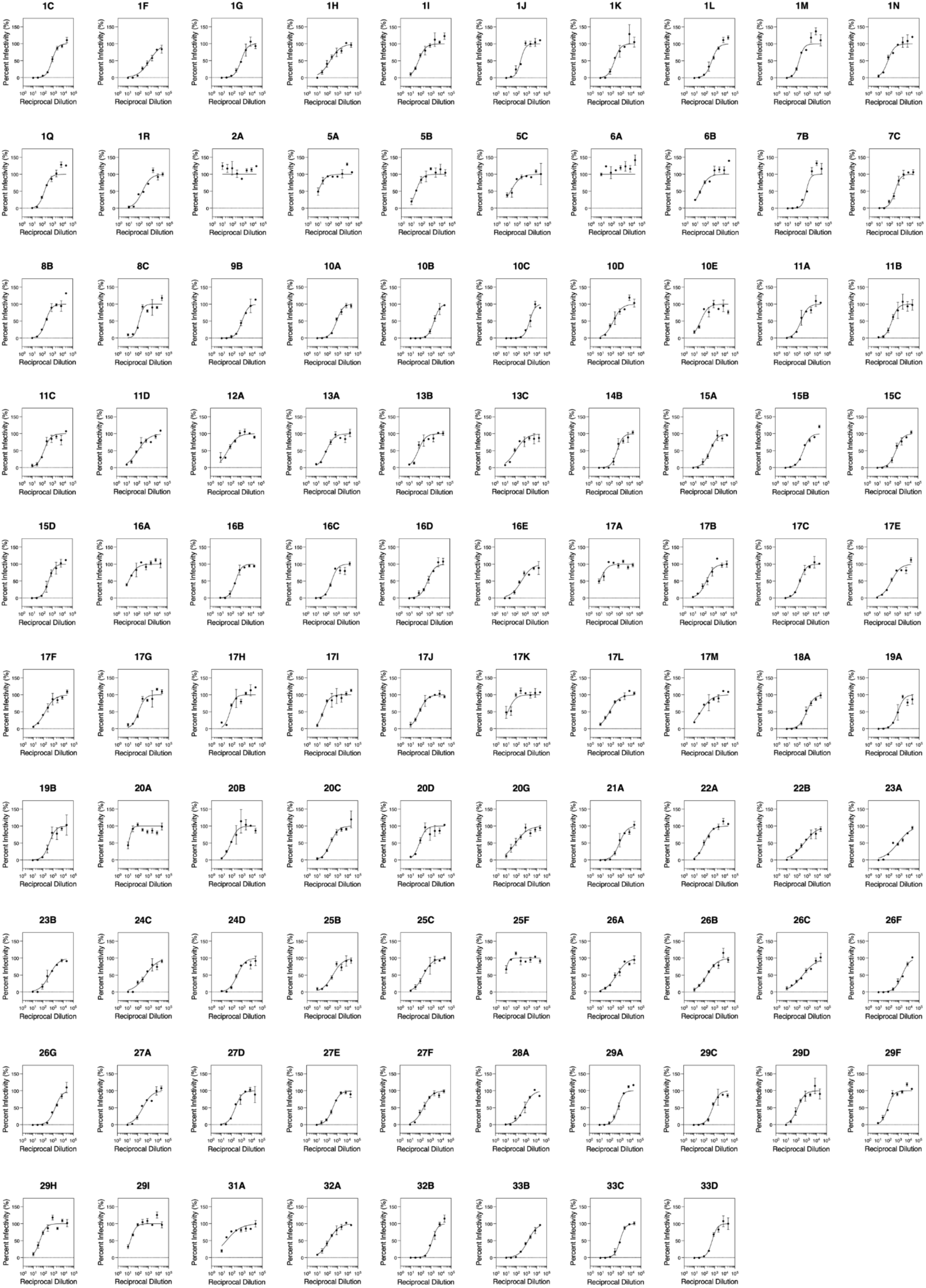
Evaluation of plasma neutralization potency against MERS-CoV EMC/2012. Dose-response curves of plasma neutralization of VSV pseudotyped with MERS-CoV EMC/2012 S for the 98 samples analyzed in this study. Data are presented as the mean ± standard error of two technical replicates from one representative biological replicate. At least two biological replicates with two technical replicates were completed for each sample using two distinct batches of pseudovirus.

**Figure S3.**
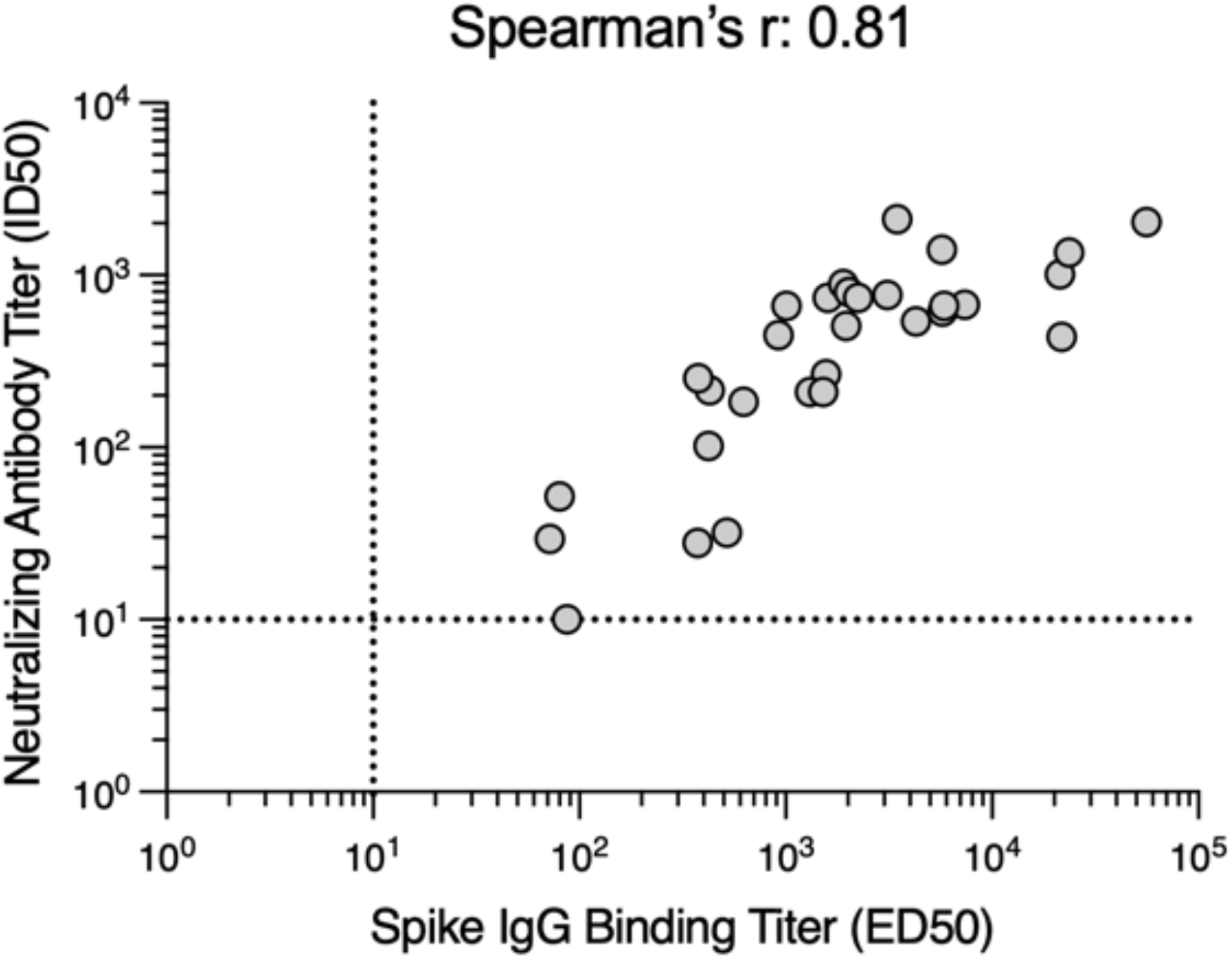
Correlation analysis of S-directed IgG binding titers and neutralizing antibody titers. The sample with highest S IgG binding titer per individual was included in the analysis. The limits of detection (ED_50_ or ID_50_: ≤ 10) are indicated with dashed lines.

**Figure S4.**
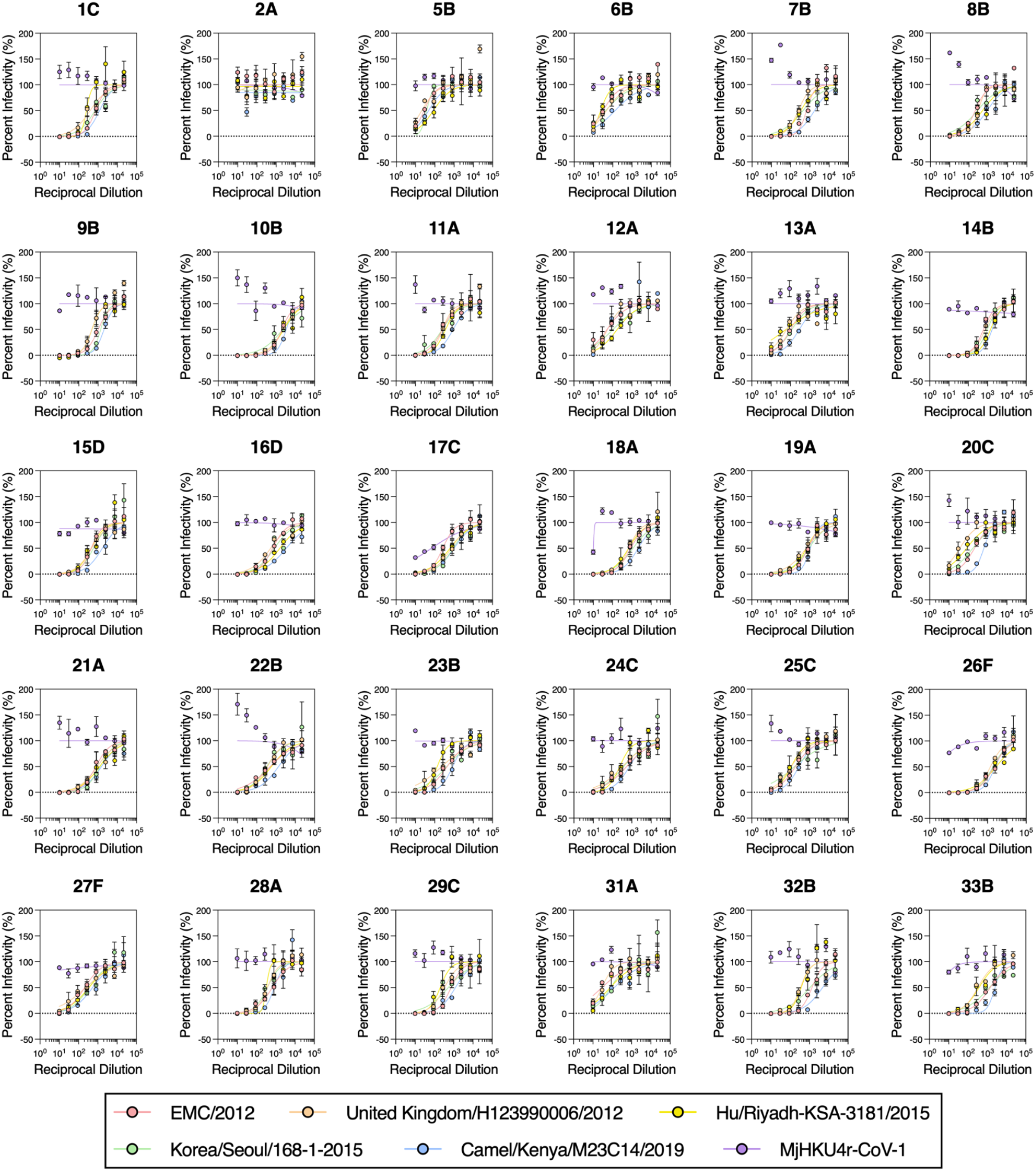
Evaluation of plasma neutralization potency against MERS-CoV variants and MjHKU4r-CoV-1. Dose-response curves for each of the 30 plasma samples included in the analysis using VSV pseudotyped with the indicated MERS-CoV variant or MjHKU4-CoV-1 S protein. Data are presented as mean ± standard error from one representative biological replicate. At least two biological replicates, each with two technical replicates, were conducted for each sample and each variant tested using unique batches of pseudovirus.

**Figure S5.**
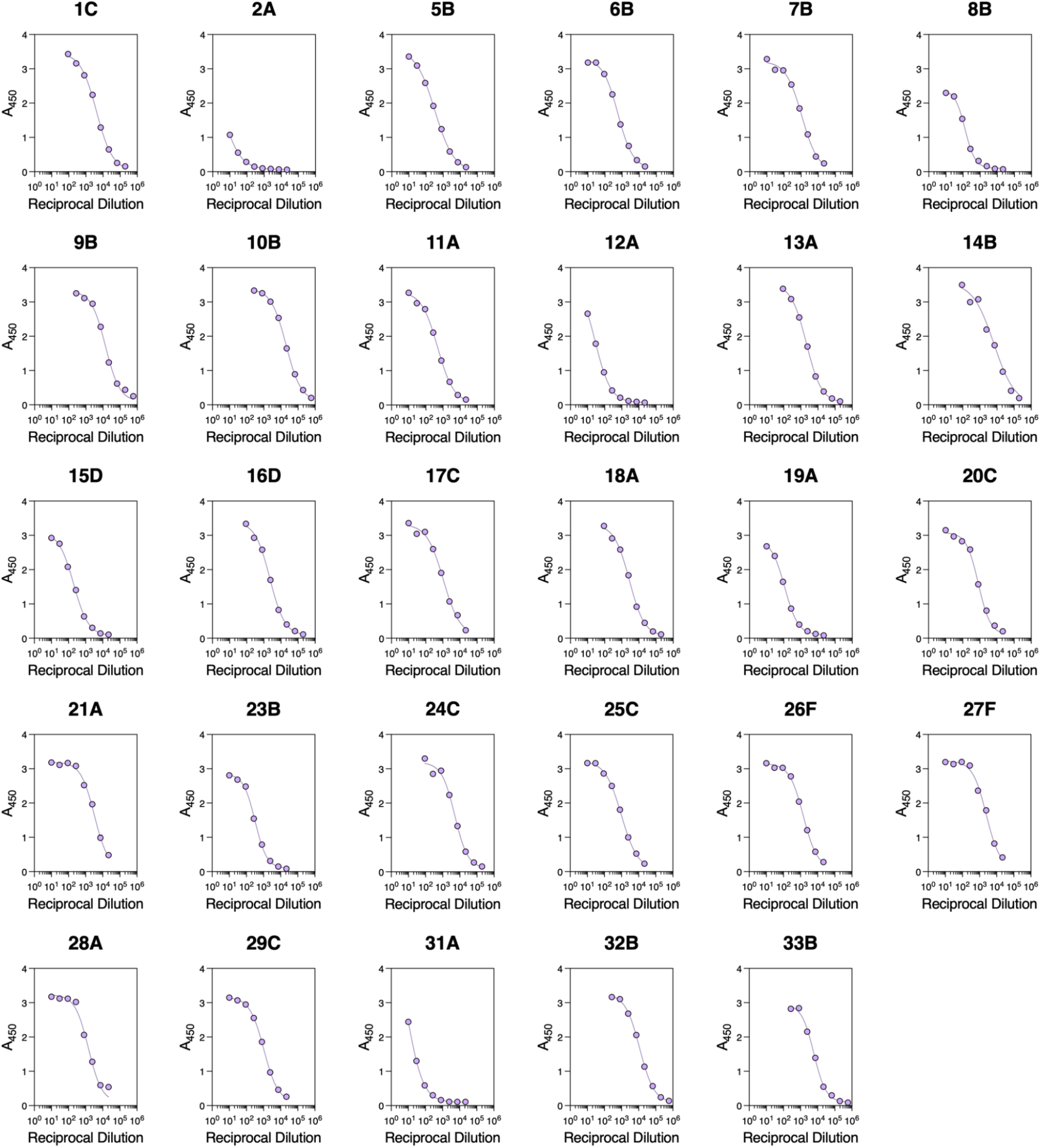
Evaluation of MERS-CoV S_1_-directed plasma IgG binding titers. Dose-response curves of plasma IgG binding to MERS-CoV S_1_ analyzed by ELISA. The sample with the highest S IgG binding titer per individual was included in the analysis. Individual 22 was excluded due to sample volume limitations. Data obtained from one representative biological replicate are presented. Two biological replicates using distinct batches of S_1_ protein were completed for each sample.

**Figure S6.**
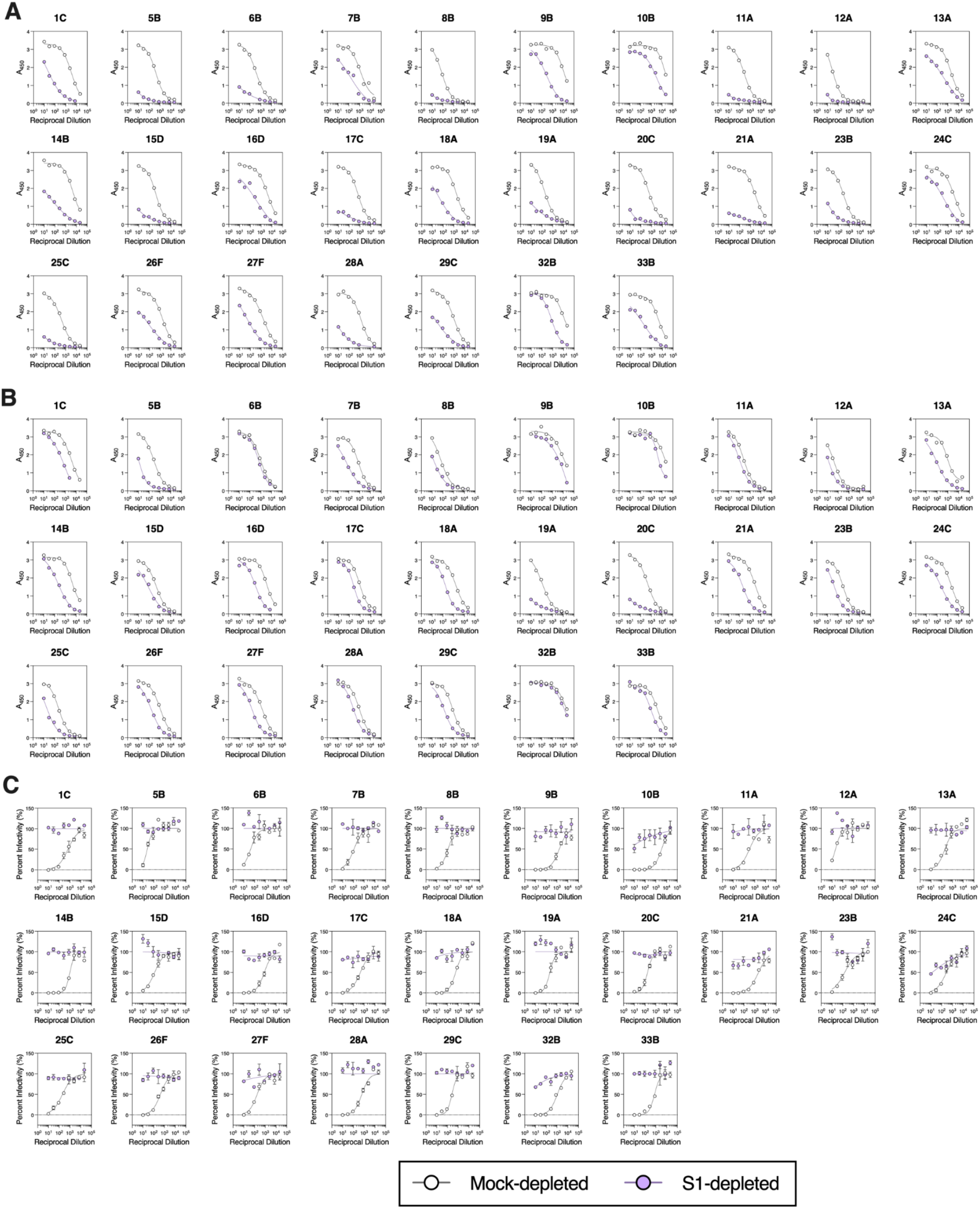
Evaluation of plasma IgG binding and neutralizing activity following depletion of MERS-CoV S_1_-directed antibodies. **A-B)** Dose-response curves of S_1_-(A) and S-directed (B) IgG binding for each of the 27 samples included in the analysis upon mock- or S_1_-depletion analyzed by ELISA using MERS-CoV EMC/2012 S_1_ or prefusion-stabilized 2P S. **C)** Dose-response curves of plasma neutralization of VSV pseudotyped with MERS-CoV EMC/2012 S for the mock- and S_1_-depleted plasma samples. Neutralization data are presented as mean ± standard error for the two technical replicates conducted. Data presented are from one representative biological replicate. Two independent biological replicates were performed using distinct batches of S_1_ protein for the antibody depletions as well as unique batches of S_1_ and S glycoproteins and pseudovirus for the ELISAs and neutralization assays, respectively.

**Figure S7.**
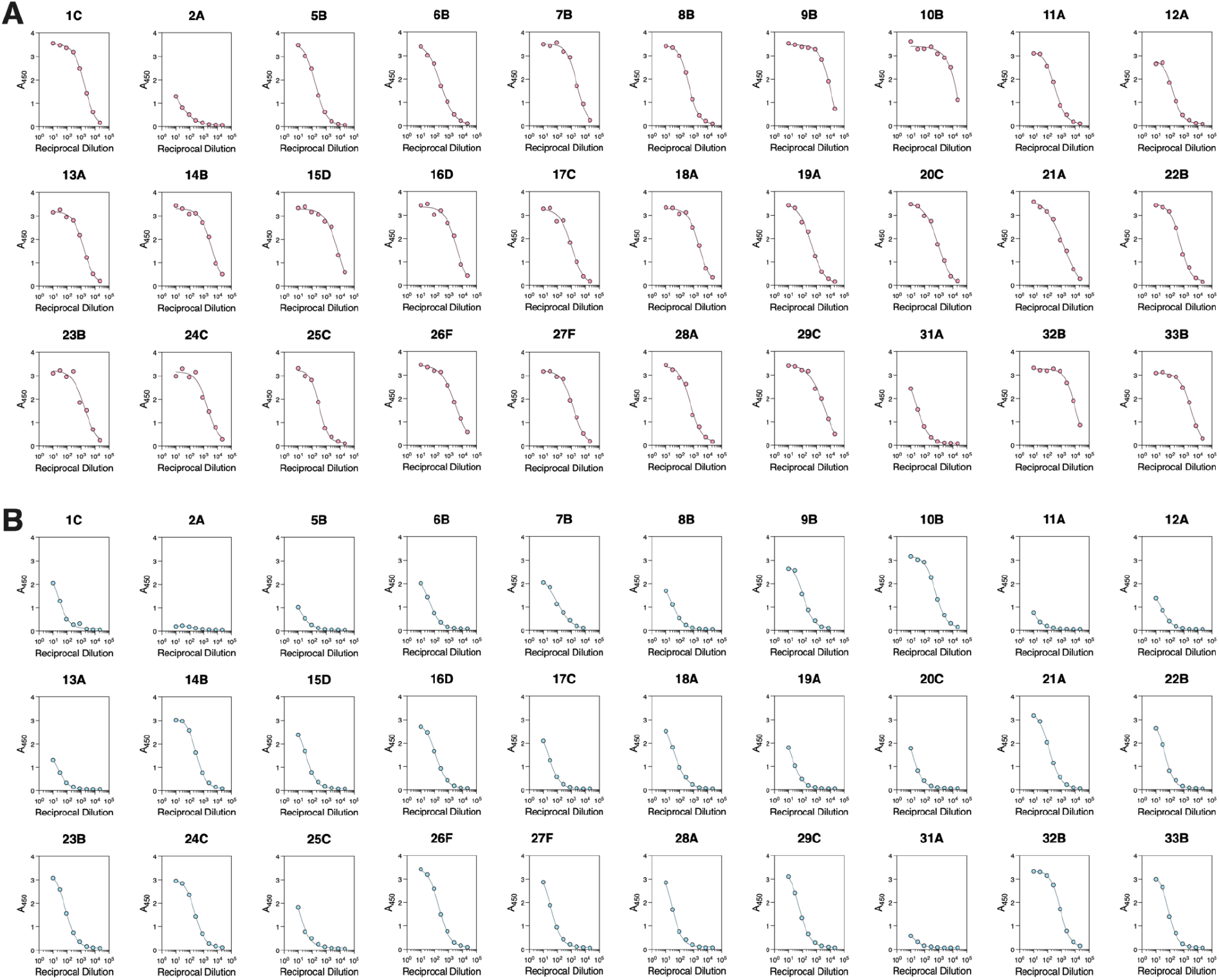
RBD and NTD IgG binding curves. **A-B)** Dose-response curves of RBD-(A) and NTD-directed (B) IgG binding for each of the 30 samples included in the analysis analyzed by ELISA using recombinantly expressed MERS-CoV EMC/2012 RBD or NTD. Data are presented from one representative biological replicate. At least two biological replicates were conducted using distinct batches of RBD or NTD.

**Figure S8.**
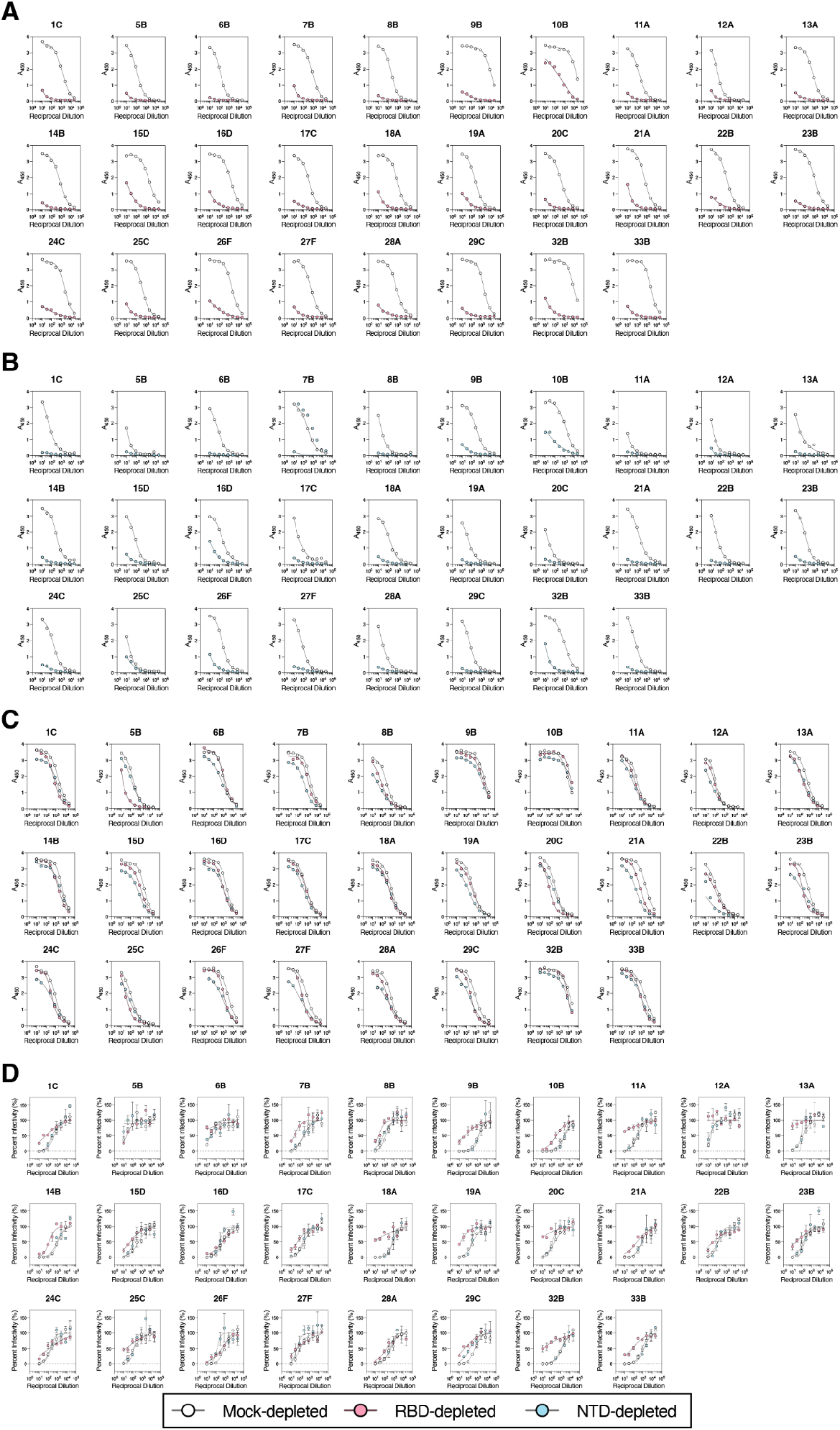
IgG binding and neutralization curves following depletion with MERS-CoV RBD and NTD. **A)** RBD IgG binding dose-response curves for each of the 28 samples included in the ELISA analysis after mock-depletion and depletion of RBD-directed antibodies. **B)** NTD IgG binding dose-response curves for each of the 28 samples included in the ELISA analysis after mock-depletion and depletion of NTD-directed antibodies. **C)** S IgG binding dose-response curves for each of the 28 samples included in the ELISA analysis after mock-depletion, depletion of RBD-directed or of NTD-directed antibodies. **D)** Dose-response curves for plasma neutralization of MERS-CoV S VSV after mock-depletion, depletion of RBD-directed or of NTD-directed antibodies. Neutralization data are presented as mean ± standard error for the two technical replicates conducted. Data presented are from one representative biological replicate. Two independent biological replicates were performed using distinct batches of RBD and NTD proteins for the antibody depletions as well as unique batches of RBD, NTD, and spike proteins and pseudovirus for the ELISAs and neutralization assays, respectively.

**Figure S9.**
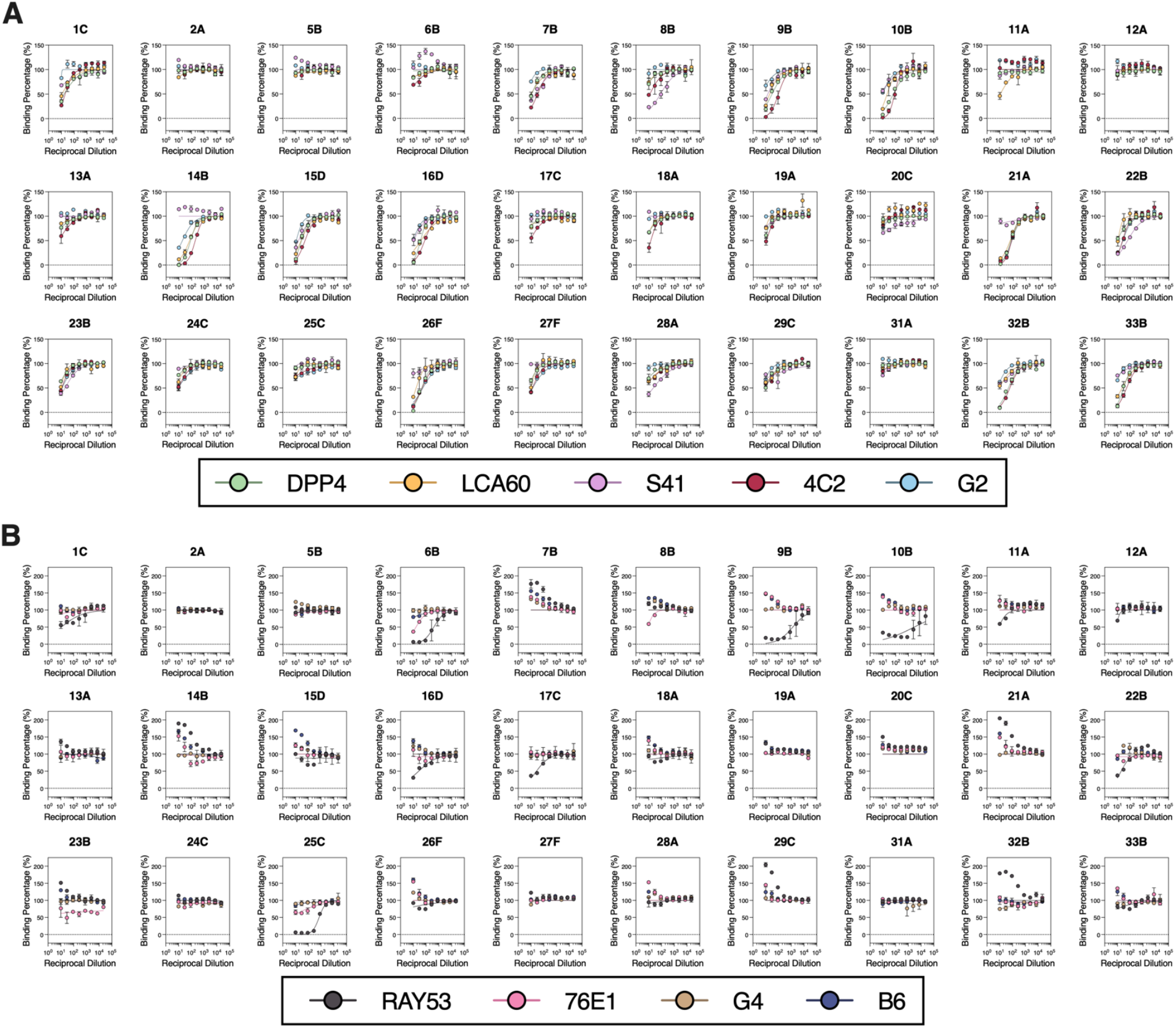
Competition ELISA curves. **A-B)** Competition ELISA curves for the 30 plasma samples analyzed using the indicated S_1_-directed (A) and S_2_-directed (B) monoclonal antibodies and prefusion-stabilized MERS-CoV EMC/2012 2P S . Data are presented as mean ± standard error for two technical replicates from one representative biological replicate. Two independent biological replicates using unique batches of biotinylated spike protein as well as monoclonal antibodies were performed for each sample.

**Table S1.** Demographic information for plasma donors.

| Individual | Sex | Age | Comorbidities |
| --- | --- | --- | --- |
| 1 | M | 42 | DM, HTN |
| 2 | M | 80 | ESRD, DM, HTN |
| 5 | M | 27 | DM, HTN |
| 6 | M | 64 | DM, HTN |
| 7 | M | 44 | DM |
| 8 | M | 51 | DM, HTN |
| 9 | M | 70 | DM ,HTN, IHD |
| 10 | M | 35 | Psychiatric Illness |
| 11 | F | 55 | DM, Hypothyroidism, AKI |
| 12 | M | 49 | DM |
| 13 | F | 41 | DM |
| 14 | M | 65 | DM, HTN, IHD |
| 15 | M | 46 | DM, HTN, COPD |
| 16 | M | 53 | DM, HTN |
| 17 | M | 49 | Heart Failure |
| 18 | M | 53 | HTN |
| 19 | M | 32 |  |
| 20 | M | 75 | DM, HTN, Epilepsy, Dyslipidemia |
| 21 | F | 67 | DM, HTN |
| 22 | M | 39 |  |
| 23 | M | 63 | DM, BA, HTN |
| 24 | M | 50 | DM |
| 25 | M | 59 | ESRD, DM, HTN |
| 26 | M | 31 | ESRD, DM, HTN |
| 27 | F | 70 | ESRD |
| 28 | F | 21 | DM |
| 29 | M | 69 | BPH, Afib |
| 31 | F | 62 | ARDS, Alzheimer, DM, HTN |
| 32 | M | 42 | DM |
| 33 | M | 58 |  |

DM: Diabetes Mellitus; HTN: Hypertension; ARDS: Acute Respiratory Distress Syndrome; ESRD: End Stage Renal Disease; IHD: Ischemic Heart Disease; AKI: Acute Kidney Injury; COPD: Chronic Obstructive Pulmonary Disease; BA: Bronchial Asthma; BPH: Benign Prostatic Hypertrophy; A fib: Atrial Fibrillation

**Table S2.** Collection timepoints for the plasma samples analyzed in this study.

| **Individual** | **Sample ID** | **Days post symptom onset or hospitalization** |
| --- | --- | --- |
| 1 | 1C | 23 |
|  | 1F | 45 |
|  | 1G | 51 |
|  | 1H | 58 |
|  | 1I | 65 |
|  | 1J | 73 |
|  | 1K | 79 |
|  | 1L | 86 |
|  | 1M | 93 |
|  | 1N | 100 |
|  | Q | 120 |
|  | 1R | 191 |
| 2 | 2A | 12 |
| 5 | 5A | 14 |
|  | 5B | 21 |
|  | 5C | 29 |
| 6 | 6A | 6 |
|  | 6B | 12 |
| 7 | 7B | 20 |
|  | 7C | 26 |
| 8 | 8B | 13 |
|  | 8C | 23 |
| 9 | 9B | 18 |
| 10 | 10A | 3 |
|  | 10B | 9 |
|  | 10C | 16 |
|  | 10D | 23 |
|  | 10E | 29 |
| 11 | 11A | 16 |
|  | 11B | 23 |
|  | 11C | 30 |
|  | 11D | 40 |
| 12 | 12A | NA |
| 13 | 13A | 12 |
|  | 13B | 19 |
|  | 13C | 26 |
| 14 | 14B | 15 |
| 15 | 15A | 12 |
|  | 15B | 19 |
|  | 15C | 26 |
|  | 15D | 33 |
| 16 | 16A | 11 |
|  | 16B | 18 |
|  | 16C | 25 |
|  | 16D | 32 |
|  | 16E | 39 |
| 17 | 17A | 5 |
|  | 17B | 12 |
|  | 17C | 19 |
|  | 17E | 33 |
|  | 17F | 40 |
|  | 17G | 47 |
|  | 17H | 54 |
|  | 17I | 61 |
|  | 17J | 68 |
|  | 17K | 74 |
|  | 17L | 82 |
|  | 17M | 89 |
| 18 | 18A | 10 |
| 19 | 19A | 6 |
|  | 19B | 15 |
| 20 | 20A | 12 |
|  | 20B | 17 |
|  | 20C | 24 |
|  | 20D | 31 |
|  | 20G | 52 |
| 21 | 21A | 7 |
| 22 | 22A | 13 |
|  | 22B | 20 |
| 23 | 23A | 7 |
|  | 23B | 14 |
| 24 | 24C | 15 |
|  | 24D | 22 |
| 25 | 25B | 81 |
|  | 25C | 88 |
|  | 25F | 109 |
| 26 | 26A | 7 |
|  | 26B | 14 |
|  | 26C | 21 |
|  | 26F | 42 |
|  | 26G | 49 |
| 27 | 27A | 10 |
|  | 27D | 31 |
|  | 27E | 38 |
|  | 27F | 45 |
| 28 | 28A | 10 |
| 29 | 29A | 10 |
|  | 29C | 24 |
|  | 29D | 31 |
|  | 29F | 45 |
|  | 29H | 59 |
|  | 29I | 66 |
| 31 | 31A | 20 |
| 32 | 32A | 5 |
|  | 32B | 12 |
| 33 | 33B | 26 |
|  | 33C | 33 |
|  | 33D | 40 |

**Table S3.** Spike protein mutations in MERS-CoV variants used in this study relative to MERS-CoV EMC/2012 (NC_019843.3).

| **MERS-CoV Variant** | **Genbank Accession Number** | **Spike Protein Mutations** |
| --- | --- | --- |
| United Kingdom/H123990006/2012 | NC_038294.1 | L506F, Q1020H |
| Hu/Riyadh-KSA-3181/2015 | KT806049.1 | L507R, Q1020R |
| Korea/Seoul/168-1-2015 | KT374056.1 | H91Y, D510G, Q1020R |
| Camel/Kenya/M23C14/2019 | OK094446.1 | V26A, D158Y, H194Y, S390F, L450F, V597A, R626P, A1158S |

## References

1. C. Drosten, S. Günther, W. Preiser, S. Van Der Werf, H.-R. Brodt, S. Becker, H. Rabenau, M. Panning, L. Kolesnikova, R. A. M. Fouchier, A. Berger, A.-M. Burguière, J. Cinatl, M. Eickmann, N. Escriou, K. Grywna, S. Kramme, J.-C. Manuguerra, S. Müller, V. Rickerts, M. Stürmer, S. Vieth, H.-D. Klenk, A. D. M. E. Osterhaus, H. Schmitz, H. W. Doerr, Identification of a Novel Coronavirus in Patients with Severe Acute Respiratory Syndrome. N Engl J Med 348, 1967–1976 (2003).

2. J. S. M. Peiris, S. T. Lai, L. L. M. Poon, Y. Guan, L. Y. C. Yam, W. Lim, J. Nicholls, W. K. S. Yee, W. W. Yan, M. T. Cheung, V. C. C. Cheng, K. H. Chan, D. N. C. Tsang, R. W. H. Yung, T. K. Ng, K. Y. Yuen, SARS study group, Coronavirus as a possible cause of severe acute respiratory syndrome. Lancet 361, 1319–1325 (2003).

3. A. M. Zaki, S. Van Boheemen, T. M. Bestebroer, A. D. M. E. Osterhaus, R. A. M. Fouchier, Isolation of a Novel Coronavirus from a Man with Pneumonia in Saudi Arabia. N Engl J Med 367, 1814–1820 (2012).

4. A. Bermingham, M. A. Chand, C. S. Brown, E. Aarons, C. Tong, C. Langrish, K. Hoschler, K. Brown, M. Galiano, R. Myers, R. G. Pebody, H. K. Green, N. L. Boddington, R. Gopal, N. Price, W. Newsholme, C. Drosten, R. A. Fouchier, M. Zambon, Severe respiratory illness caused by a novel coronavirus, in a patient transferred to the United Kingdom from the Middle East, September 2012. Eurosurveillance 17 (2012), doi:10.2807/ese.17.40.20290-en.

5. N. Zhu, D. Zhang, W. Wang, X. Li, B. Yang, J. Song, X. Zhao, B. Huang, W. Shi, R. Lu, P. Niu, F. Zhan, X. Ma, D. Wang, W. Xu, G. Wu, G. F. Gao, W. Tan, A Novel Coronavirus from Patients with Pneumonia in China, 2019. N Engl J Med 382, 727–733 (2020).

6. P. Zhou, X.-L. Yang, X.-G. Wang, B. Hu, L. Zhang, W. Zhang, H.-R. Si, Y. Zhu, B. Li, C.-L. Huang, H.-D. Chen, J. Chen, Y. Luo, H. Guo, R.-D. Jiang, M.-Q. Liu, Y. Chen, X.-R. Shen, X. Wang, X.-S. Zheng, K. Zhao, Q.-J. Chen, F. Deng, L.-L. Liu, B. Yan, F.-X. Zhan, Y.-Y. Wang, G.-F. Xiao, Z.-L. Shi, A pneumonia outbreak associated with a new coronavirus of probable bat origin. Nature 579, 270–273 (2020).

7. S. H. Ebrahim, A. D. Maher, U. Kanagasabai, S. H. Alfaraj, N. A. Alzahrani, S. A. Alqahtani, A. M. Assiri, Z. A. Memish, MERS-CoV Confirmation among 6,873 suspected persons and relevant Epidemiologic and Clinical Features, Saudi Arabia — 2014 to 2019. eClinicalMedicine 41, 101191 (2021).

8. 8. World Health Organization, Middle East Respiratory Syndrome Coronavirus - Kingdom of Saudi Arabia Disease Outbreak News (2024) (available at https://www.who.int/emergencies/disease-outbreak-news/item/2024-DON506).

9. Y. M. Arabi, A. A. Arifi, H. H. Balkhy, H. Najm, A. S. Aldawood, A. Ghabashi, H. Hawa, A. Alothman, A. Khaldi, B. Al Raiy, Clinical Course and Outcomes of Critically Ill Patients With Middle East Respiratory Syndrome Coronavirus Infection. Ann Intern Med 160, 389–397 (2014).

10. A. N. Alshukairi, J. Zheng, J. Zhao, A. Nehdi, S. A. Baharoon, L. Layqah, A. Bokhari, S. M. Al Johani, N. Samman, M. Boudjelal, P. Ten Eyck, M. A. Al-Mozaini, J. Zhao, S. Perlman, A. N. Alagaili, High Prevalence of MERS-CoV Infection in Camel Workers in Saudi Arabia. mBio 9, e01985–18 (2018).

11. G. Dudas, L. M. Carvalho, A. Rambaut, T. Bedford, MERS-CoV spillover at the camel-human interface. eLife 7, e31257 (2018).

12. V. S. Raj, H. Mou, S. L. Smits, D. H. W. Dekkers, M. A. Müller, R. Dijkman, D. Muth, J. A. A. Demmers, A. Zaki, R. A. M. Fouchier, V. Thiel, C. Drosten, P. J. M. Rottier, A. D. M. E. Osterhaus, B. J. Bosch, B. L. Haagmans, Dipeptidyl peptidase 4 is a functional receptor for the emerging human coronavirus-EMC. Nature 495, 251–254 (2013).

13. J. K. Millet, G. R. Whittaker, Host cell entry of Middle East respiratory syndrome coronavirus after two-step, furin-mediated activation of the spike protein. Proc. Natl. Acad. Sci. U.S.A. 111, 15214–15219 (2014).

14. M. A. Tortorici, D. Veesler, Structural insights into coronavirus entry. Adv Virus Res 105, 93– 116 (2019).

15. G. Lu, Y. Hu, Q. Wang, J. Qi, F. Gao, Y. Li, Y. Zhang, W. Zhang, Y. Yuan, J. Bao, B. Zhang, Y. Shi, J. Yan, G. F. Gao, Molecular basis of binding between novel human coronavirus MERS-CoV and its receptor CD26. Nature 500, 227–231 (2013).

16. N. Wang, X. Shi, L. Jiang, S. Zhang, D. Wang, P. Tong, D. Guo, L. Fu, Y. Cui, X. Liu, K. C. Arledge, Y.-H. Chen, L. Zhang, X. Wang, Structure of MERS-CoV spike receptor-binding domain complexed with human receptor DPP4. Cell Res 23, 986–993 (2013).

17. W. Li, R. J. G. Hulswit, I. Widjaja, V. S. Raj, R. McBride, W. Peng, W. Widagdo, M. A. Tortorici, B. Van Dieren, Y. Lang, J. W. M. Van Lent, J. C. Paulson, C. A. M. De Haan, R. J. De Groot, F. J. M. Van Kuppeveld, B. L. Haagmans, B.-J. Bosch, Identification of sialic acid-binding function for the Middle East respiratory syndrome coronavirus spike glycoprotein. Proc. Natl. Acad. Sci. U.S.A. 114 (2017), doi:10.1073/pnas.1712592114.

18. Y.-J. Park, A. C. Walls, Z. Wang, M. M. Sauer, W. Li, M. A. Tortorici, B.-J. Bosch, F. DiMaio, D. Veesler, Structures of MERS-CoV spike glycoprotein in complex with sialoside attachment receptors. Nat Struct Mol Biol 26, 1151–1157 (2019).

19. A. C. Walls, M. A. Tortorici, J. Snijder, X. Xiong, B.-J. Bosch, F. A. Rey, D. Veesler, Tectonic conformational changes of a coronavirus spike glycoprotein promote membrane fusion. Proc. Natl. Acad. Sci. U.S.A. 114, 11157–11162 (2017).

20. J.-E. Park, K. Li, A. Barlan, A. R. Fehr, S. Perlman, P. B. McCray, T. Gallagher, Proteolytic processing of Middle East respiratory syndrome coronavirus spikes expands virus tropism. Proc. Natl. Acad. Sci. U.S.A. 113, 12262–12267 (2016).

21. J. T. Earnest, M. P. Hantak, K. Li, P. B. McCray, S. Perlman, T. Gallagher, M. B. Frieman, Ed. The tetraspanin CD9 facilitates MERS-coronavirus entry by scaffolding host cell receptors and proteases. PLoS Pathog 13, e1006546 (2017).

22. L. Wang, W. Shi, M. G. Joyce, K. Modjarrad, Y. Zhang, K. Leung, C. R. Lees, T. Zhou, H. M. Yassine, M. Kanekiyo, Z. Yang, X. Chen, M. M. Becker, M. Freeman, L. Vogel, J. C. Johnson, G. Olinger, J. P. Todd, U. Bagci, J. Solomon, D. J. Mollura, L. Hensley, P. Jahrling, M. R. Denison, S. S. Rao, K. Subbarao, P. D. Kwong, J. R. Mascola, W.-P. Kong, B. S. Graham, Evaluation of candidate vaccine approaches for MERS-CoV. Nat Commun 6, 7712 (2015).

23. I. Widjaja, C. Wang, R. Van Haperen, J. Gutiérrez-Álvarez, B. Van Dieren, N. M. A. Okba, V. S. Raj, W. Li, R. Fernandez-Delgado, F. Grosveld, F. J. M. Van Kuppeveld, B. L. Haagmans, L. Enjuanes, D. Drabek, B.-J. Bosch, Towards a solution to MERS: protective human monoclonal antibodies targeting different domains and functions of the MERS-coronavirus spike glycoprotein. Emerging Microbes & Infections 8, 516–530 (2019).

24. J. Zhao, A. N. Alshukairi, S. A. Baharoon, W. A. Ahmed, A. A. Bokhari, A. M. Nehdi, L. A. Layqah, M. G. Alghamdi, M. M. Al Gethamy, A. M. Dada, I. Khalid, M. Boujelal, S. M. Al Johani, L. Vogel, K. Subbarao, A. Mangalam, C. Wu, P. Ten Eyck, S. Perlman, J. Zhao, Recovery from the Middle East respiratory syndrome is associated with antibody and T cell responses. Sci. Immunol. 2, eaan5393 (2017).

25. T. Koch, C. Dahlke, A. Fathi, A. Kupke, V. Krähling, N. M. A. Okba, S. Halwe, C. Rohde, M. Eickmann, A. Volz, T. Hesterkamp, A. Jambrecina, S. Borregaard, M. L. Ly, M. E. Zinser, E. Bartels, J. S. H. Poetsch, R. Neumann, R. Fux, S. Schmiedel, A. W. Lohse, B. L. Haagmans, G. Sutter, S. Becker, M. M. Addo, Safety and immunogenicity of a modified vaccinia virus Ankara vector vaccine candidate for Middle East respiratory syndrome: an open-label, phase 1 trial. Lancet Infect Dis 20, 827–838 (2020).

26. P. M. Folegatti, M. Bittaye, A. Flaxman, F. R. Lopez, D. Bellamy, A. Kupke, C. Mair, R. Makinson, J. Sheridan, C. Rohde, S. Halwe, Y. Jeong, Y.-S. Park, J.-O. Kim, M. Song, A. Boyd, N. Tran, D. Silman, I. Poulton, M. Datoo, J. Marshall, Y. Themistocleous, A. Lawrie, R. Roberts, E. Berrie, S. Becker, T. Lambe, A. Hill, K. Ewer, S. Gilbert, Safety and immunogenicity of a candidate Middle East respiratory syndrome coronavirus viral-vectored vaccine: a dose-escalation, open-label, non-randomised, uncontrolled, phase 1 trial. Lancet Infect Dis 20, 816–826 (2020).

27. K. Modjarrad, C. C. Roberts, K. T. Mills, A. R. Castellano, K. Paolino, K. Muthumani, E. L. Reuschel, M. L. Robb, T. Racine, M.-D. Oh, C. Lamarre, F. I. Zaidi, J. Boyer, S. B. Kudchodkar, M. Jeong, J. M. Darden, Y. K. Park, P. T. Scott, C. Remigio, A. P. Parikh, M. C. Wise, A. Patel, E. K. Duperret, K. Y. Kim, H. Choi, S. White, M. Bagarazzi, J. M. May, D. Kane, H. Lee, G. Kobinger, N. L. Michael, D. B. Weiner, S. J. Thomas, J. N. Maslow, Safety and immunogenicity of an anti-Middle East respiratory syndrome coronavirus DNA vaccine: a phase 1, open-label, single-arm, dose-escalation trial. Lancet Infect Dis 19, 1013–1022 (2019).

28. K. S. Corbett, D. K. Edwards, S. R. Leist, O. M. Abiona, S. Boyoglu-Barnum, R. A. Gillespie, S. Himansu, A. Schäfer, C. T. Ziwawo, A. T. DiPiazza, K. H. Dinnon, S. M. Elbashir, C. A. Shaw, A. Woods, E. J. Fritch, D. R. Martinez, K. W. Bock, M. Minai, B. M. Nagata, G. B. Hutchinson, K. Wu, C. Henry, K. Bahl, D. Garcia-Dominguez, L. Ma, I. Renzi, W.-P. Kong, S. D. Schmidt, L. Wang, Y. Zhang, E. Phung, L. A. Chang, R. J. Loomis, N. E. Altaras, E. Narayanan, M. Metkar, V. Presnyak, C. Liu, M. K. Louder, W. Shi, K. Leung, E. S. Yang, A. West, K. L. Gully, L. J. Stevens, N. Wang, D. Wrapp, N. A. Doria-Rose, G. Stewart-Jones, H. Bennett, G. S. Alvarado, M. C. Nason, T. J. Ruckwardt, J. S. McLellan, M. R. Denison, J. D. Chappell, I. N. Moore, K. M. Morabito, J. R. Mascola, R. S. Baric, A. Carfi, B. S. Graham, SARS-CoV-2 mRNA vaccine design enabled by prototype pathogen preparedness. Nature 586, 567–571 (2020).

29. D. Corti, J. Zhao, M. Pedotti, L. Simonelli, S. Agnihothram, C. Fett, B. Fernandez-Rodriguez, M. Foglierini, G. Agatic, F. Vanzetta, R. Gopal, C. J. Langrish, N. A. Barrett, F. Sallusto, R. S. Baric, L. Varani, M. Zambon, S. Perlman, A. Lanzavecchia, Prophylactic and postexposure efficacy of a potent human monoclonal antibody against MERS coronavirus. Proc. Natl. Acad. Sci. U.S.A. 112, 10473–10478 (2015).

30. A. C. Walls, X. Xiong, Y.-J. Park, M. A. Tortorici, J. Snijder, J. Quispe, E. Cameroni, R. Gopal, M. Dai, A. Lanzavecchia, M. Zambon, F. A. Rey, D. Corti, D. Veesler, Unexpected Receptor Functional Mimicry Elucidates Activation of Coronavirus Fusion. Cell 176, 1026–1039.e15 (2019).

31. S. Zhang, W. Jia, J. Zeng, M. Li, Z. Wang, H. Zhou, L. Zhang, X. Wang, Cryoelectron microscopy structures of a human neutralizing antibody bound to MERS-CoV spike glycoprotein. Front. Microbiol. 13, 988298 (2022).

32. S. Zhang, P. Zhou, P. Wang, Y. Li, L. Jiang, W. Jia, H. Wang, A. Fan, D. Wang, X. Shi, X. Fang, M. Hammel, S. Wang, X. Wang, L. Zhang, Structural Definition of a Unique Neutralization Epitope on the Receptor-Binding Domain of MERS-CoV Spike Glycoprotein. Cell Reports 24, 441–452 (2018).

33. L. Jiang, N. Wang, T. Zuo, X. Shi, K.-M. V. Poon, Y. Wu, F. Gao, D. Li, R. Wang, J. Guo, L. Fu, K.-Y. Yuen, B.-J. Zheng, X. Wang, L. Zhang, Potent Neutralization of MERS-CoV by Human Neutralizing Monoclonal Antibodies to the Viral Spike Glycoprotein. Sci. Transl. Med. 6 (2014), doi:10.1126/scitranslmed.3008140.

34. Y. Li, Y. Wan, P. Liu, J. Zhao, G. Lu, J. Qi, Q. Wang, X. Lu, Y. Wu, W. Liu, B. Zhang, K.-Y. Yuen, S. Perlman, G. F. Gao, J. Yan, A humanized neutralizing antibody against MERS-CoV targeting the receptor-binding domain of the spike protein. Cell Res 25, 1237–1249 (2015).

35. H. Zhou, Y. Chen, S. Zhang, P. Niu, K. Qin, W. Jia, B. Huang, S. Zhang, J. Lan, L. Zhang, W. Tan, X. Wang, Structural definition of a neutralization epitope on the N-terminal domain of MERS-CoV spike glycoprotein. Nat Commun 10, 3068 (2019).

36. J. Pallesen, N. Wang, K. S. Corbett, D. Wrapp, R. N. Kirchdoerfer, H. L. Turner, C. A. Cottrell, M. M. Becker, L. Wang, W. Shi, W.-P. Kong, E. L. Andres, A. N. Kettenbach, M. R. Denison, J. D. Chappell, B. S. Graham, A. B. Ward, J. S. McLellan, Immunogenicity and structures of a rationally designed prefusion MERS-CoV spike antigen. Proc. Natl. Acad. Sci. U.S.A. 114 (2017), doi:10.1073/pnas.1707304114.

37. X.-C. Tang, S. S. Agnihothram, Y. Jiao, J. Stanhope, R. L. Graham, E. C. Peterson, Y. Avnir, A. St. C. Tallarico, J. Sheehan, Q. Zhu, R. S. Baric, W. A. Marasco, Identification of human neutralizing antibodies against MERS-CoV and their role in virus adaptive evolution. Proc. Natl. Acad. Sci. U.S.A. 111 (2014), doi:10.1073/pnas.1402074111.

38. X. Sun, C. Yi, Y. Zhu, L. Ding, S. Xia, X. Chen, M. Liu, C. Gu, X. Lu, Y. Fu, S. Chen, T. Zhang, Y. Zhang, Z. Yang, L. Ma, W. Gu, G. Hu, S. Du, R. Yan, W. Fu, S. Yuan, C. Qiu, C. Zhao, X. Zhang, Y. He, A. Qu, X. Zhou, X. Li, G. Wong, Q. Deng, Q. Zhou, H. Lu, Z. Ling, J. Ding, L. Lu, J. Xu, Y. Xie, B. Sun, Neutralization mechanism of a human antibody with pan-coronavirus reactivity including SARS-CoV-2. Nat Microbiol 7, 1063–1074 (2022).

39. R. P. Silva, Y. Huang, A. W. Nguyen, C.-L. Hsieh, O. S. Olaluwoye, T. S. Kaoud, R. E. Wilen, A. N. Qerqez, J.-G. Park, A. M. Khalil, L. R. Azouz, K. C. Le, A. L. Bohanon, A. M. DiVenere, Y. Liu, A. G. Lee, D. A. Amengor, S. R. Shoemaker, S. M. Costello, E. A. Padlan, S. Marqusee, L. Martinez-Sobrido, K. N. Dalby, S. D’Arcy, J. S. McLellan, J. A. Maynard, Identification of a conserved S2 epitope present on spike proteins from all highly pathogenic coronaviruses. eLife 12, e83710 (2023).

40. M. M. Sauer, M. A. Tortorici, Y.-J. Park, A. C. Walls, L. Homad, O. J. Acton, J. E. Bowen, C. Wang, X. Xiong, W. De Van Der Schueren, J. Quispe, B. G. Hoffstrom, B.-J. Bosch, A. T. McGuire, D. Veesler, Structural basis for broad coronavirus neutralization. Nat Struct Mol Biol 28, 478– 486 (2021).

41. D. Pinto, M. M. Sauer, N. Czudnochowski, J. S. Low, M. A. Tortorici, M. P. Housley, J. Noack, A. C. Walls, J. E. Bowen, B. Guarino, L. E. Rosen, J. Di Iulio, J. Jerak, H. Kaiser, S. Islam, S. Jaconi, N. Sprugasci, K. Culap, R. Abdelnabi, C. Foo, L. Coelmont, I. Bartha, S. Bianchi, C. Silacci-Fregni, J. Bassi, R. Marzi, E. Vetti, A. Cassotta, A. Ceschi, P. Ferrari, P. E. Cippà, O. Giannini, S. Ceruti, C. Garzoni, A. Riva, F. Benigni, E. Cameroni, L. Piccoli, M. S. Pizzuto, M. Smithey, D. Hong, A. Telenti, F. A. Lempp, J. Neyts, C. Havenar-Daughton, A. Lanzavecchia, F. Sallusto, G. Snell, H. W. Virgin, M. Beltramello, D. Corti, D. Veesler, Broad betacoronavirus neutralization by a stem helix–specific human antibody. Science 373, 1109–1116 (2021).

42. L. V. Tse, Y. J. Hou, E. McFadden, R. E. Lee, T. D. Scobey, S. R. Leist, D. R. Martinez, R. M. Meganck, A. Schäfer, B. L. Yount, T. Mascenik, J. M. Powers, S. H. Randell, Y. Zhang, L. Wang, J. Mascola, J. S. McLellan, R. S. Baric, A MERS-CoV antibody neutralizes a pre-emerging group 2c bat coronavirus. Sci. Transl. Med. 15, eadg5567 (2023).

43. N. Wang, O. Rosen, L. Wang, H. L. Turner, L. J. Stevens, K. S. Corbett, C. A. Bowman, J. Pallesen, W. Shi, Y. Zhang, K. Leung, R. N. Kirchdoerfer, M. M. Becker, M. R. Denison, J. D. Chappell, A. B. Ward, B. S. Graham, J. S. McLellan, Structural Definition of a Neutralization-Sensitive Epitope on the MERS-CoV S1-NTD. Cell Rep 28, 3395–3405.e6 (2019).

44. L. Piccoli, Y.-J. Park, M. A. Tortorici, N. Czudnochowski, A. C. Walls, M. Beltramello, C. Silacci-Fregni, D. Pinto, L. E. Rosen, J. E. Bowen, O. J. Acton, S. Jaconi, B. Guarino, A. Minola, F. Zatta, N. Sprugasci, J. Bassi, A. Peter, A. De Marco, J. C. Nix, F. Mele, S. Jovic, B. F. Rodriguez, S. V. Gupta, F. Jin, G. Piumatti, G. Lo Presti, A. F. Pellanda, M. Biggiogero, M. Tarkowski, M. S. Pizzuto, E. Cameroni, C. Havenar-Daughton, M. Smithey, D. Hong, V. Lepori, E. Albanese, A. Ceschi, E. Bernasconi, L. Elzi, P. Ferrari, C. Garzoni, A. Riva, G. Snell, F. Sallusto, K. Fink, H. W. Virgin, A. Lanzavecchia, D. Corti, D. Veesler, Mapping Neutralizing and Immunodominant Sites on the SARS-CoV-2 Spike Receptor-Binding Domain by Structure-Guided High-Resolution Serology. Cell 183, 1024–1042.e21 (2020).

45. L. Premkumar, B. Segovia-Chumbez, R. Jadi, D. R. Martinez, R. Raut, A. J. Markmann, C. Cornaby, L. Bartelt, S. Weiss, Y. Park, C. E. Edwards, E. Weimer, E. M. Scherer, N. Rouphael, S. Edupuganti, D. Weiskopf, L. V. Tse, Y. J. Hou, D. Margolis, A. Sette, M. H. Collins, J. Schmitz, R. S. Baric, A. M. De Silva, The receptor-binding domain of the viral spike protein is an immunodominant and highly specific target of antibodies in SARS-CoV-2 patients. Sci. Immunol. 5, eabc8413 (2020).

46. J. E. Bowen, Y.-J. Park, C. Stewart, J. T. Brown, W. K. Sharkey, A. C. Walls, A. Joshi, K. R. Sprouse, M. McCallum, M. A. Tortorici, N. M. Franko, J. K. Logue, I. G. Mazzitelli, A. W. Nguyen, R. P. Silva, Y. Huang, J. S. Low, J. Jerak, S. W. Tiles, K. Ahmed, A. Shariq, J. M. Dan, Z. Zhang, D. Weiskopf, A. Sette, G. Snell, C. M. Posavad, N. T. Iqbal, J. Geffner, A. Bandera, A. Gori, F. Sallusto, J. A. Maynard, S. Crotty, W. C. Van Voorhis, C. Simmerling, R. Grifantini, H. Y. Chu, D. Corti, D. Veesler, SARS-CoV-2 spike conformation determines plasma neutralizing activity elicited by a wide panel of human vaccines. *Sci. Immunol.*, eadf1421 (2022).

47. A. J. Greaney, A. N. Loes, K. H. D. Crawford, T. N. Starr, K. D. Malone, H. Y. Chu, J. D. Bloom, Comprehensive mapping of mutations in the SARS-CoV-2 receptor-binding domain that affect recognition by polyclonal human plasma antibodies. Cell Host & Microbe 29, 463–476.e6 (2021).

48. A. J. Greaney, R. T. Eguia, T. N. Starr, K. Khan, N. Franko, J. K. Logue, S. M. Lord, C. Speake, H. Y. Chu, A. Sigal, J. D. Bloom, M. Dittmann, Ed. The SARS-CoV-2 Delta variant induces an antibody response largely focused on class 1 and 2 antibody epitopes. PLoS Pathog 18, e1010592 (2022).

49. A. J. Greaney, T. N. Starr, R. T. Eguia, A. N. Loes, K. Khan, F. Karim, S. Cele, J. E. Bowen, J. K. Logue, D. Corti, D. Veesler, H. Y. Chu, A. Sigal, J. D. Bloom, S. L. Klein, Ed. A SARS-CoV-2 variant elicits an antibody response with a shifted immunodominance hierarchy. PLoS Pathog 18, e1010248 (2022).

50. A. N. Alshukairi, J. Zhao, M. A. Al-Mozaini, Y. Wang, A. Dada, S. A. Baharoon, S. Alfaraj, W. A. Ahmed, M. A. Enani, F. E. Elzein, N. Eltayeb, L. Layqah, A. El-Saed, H. A. Bahaudden, A. Haseeb, S. A. El-Kafrawy, A. M. Hassan, N. A. Siddiq, I. Alsharif, I. Qushmaq, E. I. Azhar, S. Perlman, Z. A. Memish, Longevity of Middle East Respiratory Syndrome Coronavirus Antibody Responses in Humans, Saudi Arabia. Emerg. Infect. Dis. 27 (2021), doi:10.3201/eid2705.204056.

51. S. Cheon, U. Park, H. Park, Y. Kim, Y. T. H. Nguyen, A. Aigerim, J.-Y. Rhee, J.-P. Choi, W. B. Park, S. W. Park, Y. Kim, D.-G. Lim, J.-S. Yang, J.-Y. Lee, Y.-S. Kim, N.-H. Cho, Longevity of seropositivity and neutralizing antibodies in recovered MERS patients: a 5-year follow-up study. Clinical Microbiology and Infection 28, 292–296 (2022).

52. P. G. Choe, R. a. P. M. Perera, W. B. Park, K.-H. Song, J. H. Bang, E. S. Kim, H. B. Kim, L. W. R. Ko, S. W. Park, N.-J. Kim, E. H. Y. Lau, L. L. M. Poon, M. Peiris, M.-D. Oh, MERS-CoV Antibody Responses 1 Year after Symptom Onset, South Korea, 2015. Emerg Infect Dis 23, 1079–1084 (2017).

53. W. B. Park, R. A. P. M. Perera, P. G. Choe, E. H. Y. Lau, S. J. Choi, J. Y. Chun, H. S. Oh, K.-H. Song, J. H. Bang, E. S. Kim, H. B. Kim, S. W. Park, N. J. Kim, L. L. Man Poon, M. Peiris, M.-D. Oh, Kinetics of Serologic Responses to MERS Coronavirus Infection in Humans, South Korea. Emerg Infect Dis 21, 2186–2189 (2015).

54. D. C. Payne, I. Iblan, B. Rha, S. Alqasrawi, A. Haddadin, M. Al Nsour, T. Alsanouri, S. S. Ali, J. Harcourt, C. Miao, A. Tamin, S. I. Gerber, L. M. Haynes, M. M. Al Abdallat, Persistence of Antibodies against Middle East Respiratory Syndrome Coronavirus. Emerg Infect Dis 22, 1824– 1826 (2016).

55. I. Ngere, E. A. Hunsperger, S. Tong, J. Oyugi, W. Jaoko, J. L. Harcourt, N. J. Thornburg, H. Oyas, M. Muturi, E. M. Osoro, J. Gachohi, C. Ombok, J. Dawa, Y. Tao, J. Zhang, L. Mwasi, C. Ochieng, A. Mwatondo, B. Bodha, D. Langat, A. Herman-Roloff, M. K. Njenga, M.-A. Widdowson, P. M. Munyua, Outbreak of Middle East Respiratory Syndrome Coronavirus in Camels and Probable Spillover Infection to Humans in Kenya. Viruses 14, 1743 (2022).

56. W. Tai, Y. Wang, C. A. Fett, G. Zhao, F. Li, S. Perlman, S. Jiang, Y. Zhou, L. Du, Recombinant Receptor-Binding Domains of Multiple Middle East Respiratory Syndrome Coronaviruses (MERS-CoVs) Induce Cross-Neutralizing Antibodies against Divergent Human and Camel MERS-CoVs and Antibody Escape Mutants. J Virol 91, e01651–16 (2017).

57. T.-H. Jang, W.-J. Park, H. Lee, H.-M. Woo, S.-Y. Lee, K.-C. Kim, S. S. Kim, E. Hong, J. Song, J.-Y. Lee, The structure of a novel antibody against the spike protein inhibits Middle East respiratory syndrome coronavirus infections. Sci Rep 12, 1260 (2022).

58. H. Kleine-Weber, M. T. Elzayat, L. Wang, B. S. Graham, M. A. Müller, C. Drosten, S. Pöhlmann, M. Hoffmann, Mutations in the Spike Protein of Middle East Respiratory Syndrome Coronavirus Transmitted in Korea Increase Resistance to Antibody-Mediated Neutralization. J Virol 93, e01381–18 (2019).

59. Y. Kim, S. Cheon, C.-K. Min, K. M. Sohn, Y. J. Kang, Y.-J. Cha, J.-I. Kang, S. K. Han, N.-Y. Ha, G. Kim, A. Aigerim, H. M. Shin, M.-S. Choi, S. Kim, H.-S. Cho, Y.-S. Kim, N.-H. Cho, M. J. Buchmeier, Ed. Spread of Mutant Middle East Respiratory Syndrome Coronavirus with Reduced Affinity to Human CD26 during the South Korean Outbreak. mBio 7, e00019–16 (2016).

60. L.-Y. R. Wong, J. Zheng, A. Sariol, S. Lowery, D. K. Meyerholz, T. Gallagher, S. Perlman, Middle East respiratory syndrome coronavirus Spike protein variants exhibit geographic differences in virulence. Proc. Natl. Acad. Sci. U.S.A. 118, e2102983118 (2021).

61. J. Chen, X. Yang, H. Si, Q. Gong, T. Que, J. Li, Y. Li, C. Wu, W. Zhang, Y. Chen, Y. Luo, Y. Zhu, B. Li, D. Luo, B. Hu, H. Lin, R. Jiang, T. Jiang, Q. Li, M. Liu, S. Xie, J. Su, X. Zheng, A. Li, Y. Yao, Y. Yang, P. Chen, A. Wu, M. He, X. Lin, Y. Tong, Y. Hu, Z.-L. Shi, P. Zhou, A bat MERS-like coronavirus circulates in pangolins and utilizes human DPP4 and host proteases for cell entry. Cell 186, 850–863.e16 (2023).

62. M. A. Tortorici, M. Beltramello, F. A. Lempp, D. Pinto, H. V. Dang, L. E. Rosen, M. McCallum, J. Bowen, A. Minola, S. Jaconi, F. Zatta, A. De Marco, B. Guarino, S. Bianchi, E. J. Lauron, H. Tucker, J. Zhou, A. Peter, C. Havenar-Daughton, J. A. Wojcechowskyj, J. B. Case, R. E. Chen, H. Kaiser, M. Montiel-Ruiz, M. Meury, N. Czudnochowski, R. Spreafico, J. Dillen, C. Ng, N. Sprugasci, K. Culap, F. Benigni, R. Abdelnabi, S.-Y. C. Foo, M. A. Schmid, E. Cameroni, A. Riva, A. Gabrieli, M. Galli, M. S. Pizzuto, J. Neyts, M. S. Diamond, H. W. Virgin, G. Snell, D. Corti, K. Fink, D. Veesler, Ultrapotent human antibodies protect against SARS-CoV-2 challenge via multiple mechanisms. Science 370, 950–957 (2020).

63. K. H. D. Crawford, A. S. Dingens, R. Eguia, C. R. Wolf, N. Wilcox, J. K. Logue, K. Shuey, A. M. Casto, B. Fiala, S. Wrenn, D. Pettie, N. P. King, A. L. Greninger, H. Y. Chu, J. D. Bloom, Dynamics of Neutralizing Antibody Titers in the Months After Severe Acute Respiratory Syndrome Coronavirus 2 Infection. J Infect Dis 223, 197–205 (2021).

64. M. McCallum, J. Bassi, A. De Marco, A. Chen, A. C. Walls, J. Di Iulio, M. A. Tortorici, M.-J. Navarro, C. Silacci-Fregni, C. Saliba, K. R. Sprouse, M. Agostini, D. Pinto, K. Culap, S. Bianchi, S. Jaconi, E. Cameroni, J. E. Bowen, S. W. Tilles, M. S. Pizzuto, S. B. Guastalla, G. Bona, A. F. Pellanda, C. Garzoni, W. C. Van Voorhis, L. E. Rosen, G. Snell, A. Telenti, H. W. Virgin, L. Piccoli, D. Corti, D. Veesler, SARS-CoV-2 immune evasion by the B.1.427/B.1.429 variant of concern. Science, eabi7994 (2021).

65. M. McCallum, A. C. Walls, K. R. Sprouse, J. E. Bowen, L. E. Rosen, H. V. Dang, A. De Marco, N. Franko, S. W. Tilles, J. Logue, M. C. Miranda, M. Ahlrichs, L. Carter, G. Snell, M. S. Pizzuto, H. Y. Chu, W. C. Van Voorhis, D. Corti, D. Veesler, Molecular basis of immune evasion by the Delta and Kappa SARS-CoV-2 variants. Science 374, 1621–1626 (2021).

66. M. McCallum, A. De Marco, F. A. Lempp, M. A. Tortorici, D. Pinto, A. C. Walls, M. Beltramello, A. Chen, Z. Liu, F. Zatta, S. Zepeda, J. di Iulio, J. E. Bowen, M. Montiel-Ruiz, J. Zhou, L. E. Rosen, S. Bianchi, B. Guarino, C. S. Fregni, R. Abdelnabi, S.-Y. C. Foo, P. W. Rothlauf, L.-M. Bloyet, F. Benigni, E. Cameroni, J. Neyts, A. Riva, G. Snell, A. Telenti, S. P. J. Whelan, H. W. Virgin, D. Corti, M. S. Pizzuto, D. Veesler, N-terminal domain antigenic mapping reveals a site of vulnerability for SARS-CoV-2. Cell 184, 2332–2347.e16 (2021).

67. A. C. Walls, B. Fiala, A. Schäfer, S. Wrenn, M. N. Pham, M. Murphy, L. V. Tse, L. Shehata, M. A. O’Connor, C. Chen, M. J. Navarro, M. C. Miranda, D. Pettie, R. Ravichandran, J. C. Kraft, C. Ogohara, A. Palser, S. Chalk, E.-C. Lee, K. Guerriero, E. Kepl, C. M. Chow, C. Sydeman, E. A. Hodge, B. Brown, J. T. Fuller, K. H. Dinnon, L. E. Gralinski, S. R. Leist, K. L. Gully, T. B. Lewis, M. Guttman, H. Y. Chu, K. K. Lee, D. H. Fuller, R. S. Baric, P. Kellam, L. Carter, M. Pepper, T. P. Sheahan, D. Veesler, N. P. King, Elicitation of Potent Neutralizing Antibody Responses by Designed Protein Nanoparticle Vaccines for SARS-CoV-2. Cell 183, 1367–1382.e17 (2020).

68. A. C. Walls, M. C. Miranda, A. Schäfer, M. N. Pham, A. Greaney, P. S. Arunachalam, M.-J. Navarro, M. A. Tortorici, K. Rogers, M. A. O’Connor, L. Shirreff, D. E. Ferrell, J. Bowen, N. Brunette, E. Kepl, S. K. Zepeda, T. Starr, C.-L. Hsieh, B. Fiala, S. Wrenn, D. Pettie, C. Sydeman, K. R. Sprouse, M. Johnson, A. Blackstone, R. Ravichandran, C. Ogohara, L. Carter, S. W. Tilles, R. Rappuoli, S. R. Leist, D. R. Martinez, M. Clark, R. Tisch, D. T. O’Hagan, R. Van Der Most, W. C. Van Voorhis, D. Corti, J. S. McLellan, H. Kleanthous, T. P. Sheahan, K. D. Smith, D. H. Fuller, F. Villinger, J. Bloom, B. Pulendran, R. S. Baric, N. P. King, D. Veesler, Elicitation of broadly protective sarbecovirus immunity by receptor-binding domain nanoparticle vaccines. Cell 184, 5432–5447.e16 (2021).

69. T. N. Starr, N. Czudnochowski, Z. Liu, F. Zatta, Y.-J. Park, A. Addetia, D. Pinto, M. Beltramello, P. Hernandez, A. J. Greaney, R. Marzi, W. G. Glass, I. Zhang, A. S. Dingens, J. E. Bowen, M. A. Tortorici, A. C. Walls, J. A. Wojcechowskyj, A. De Marco, L. E. Rosen, J. Zhou, M. Montiel-Ruiz, H. Kaiser, J. Dillen, H. Tucker, J. Bassi, C. Silacci-Fregni, M. P. Housley, J. di Iulio, G. Lombardo, M. Agostini, N. Sprugasci, K. Culap, S. Jaconi, M. Meury, E. Dellota, R. Abdelnabi, S.-Y. C. Foo, E. Cameroni, S. Stumpf, T. I. Croll, J. C. Nix, C. Havenar-Daughton, L. Piccoli, F. Benigni, J. Neyts, A. Telenti, F. A. Lempp, M. S. Pizzuto, J. D. Chodera, C. M. Hebner, H. W. Virgin, S. P. J. Whelan, D. Veesler, D. Corti, J. D. Bloom, G. Snell, SARS-CoV-2 RBD antibodies that maximize breadth and resistance to escape. Nature (2021), doi:10.1038/s41586-021-03807-6.

70. C. O. Barnes, C. A. Jette, M. E. Abernathy, K.-M. A. Dam, S. R. Esswein, H. B. Gristick, A. G. Malyutin, N. G. Sharaf, K. E. Huey-Tubman, Y. E. Lee, D. F. Robbiani, M. C. Nussenzweig, A. P. West, P. J. Bjorkman, SARS-CoV-2 neutralizing antibody structures inform therapeutic strategies. Nature 588, 682–687 (2020).

71. J. S. Low, J. Jerak, M. A. Tortorici, M. McCallum, D. Pinto, A. Cassotta, M. Foglierini, F. Mele, R. Abdelnabi, B. Weynand, J. Noack, M. Montiel-Ruiz, S. Bianchi, F. Benigni, N. Sprugasci, A. Joshi, J. E. Bowen, C. Stewart, M. Rexhepaj, A. C. Walls, D. Jarrossay, D. Morone, P. Paparoditis, C. Garzoni, P. Ferrari, A. Ceschi, J. Neyts, L. A. Purcell, G. Snell, D. Corti, A. Lanzavecchia, D. Veesler, F. Sallusto, ACE2-binding exposes the SARS-CoV-2 fusion peptide to broadly neutralizing coronavirus antibodies. Science 377, 735–742 (2022).

72. 72. J. Lee, C. Stewart, A. Schaefer, E. M. Leaf, Y.-J. Park, D. Asarnow, J. M. Powers, C. Treichel, D. Corti, R. Baric, N. P. King, D. Veesler, A broadly generalizable stabilization strategy for sarbecovirus fusion machinery vaccines (Biochemistry, 2023; http://biorxiv.org/lookup/doi/10.1101/2023.12.12.571160).

73. 73. X. Nuqui, L. Casalino, L. Zhou, M. Shehata, A. Wang, A. L. Tse, A. A. Ojha, F. L. Kearns, M. A. Rosenfeld, E. H. Miller, C. M. Acreman, S.-H. Ahn, K. Chandran, J. S. McLellan, R. E. Amaro, Simulation-Driven Design of Stabilized SARS-CoV-2 Spike S2 Immunogens (Biophysics, 2023; http://biorxiv.org/lookup/doi/10.1101/2023.10.24.563841).

74. C.-L. Hsieh, A. P. Werner, S. R. Leist, L. J. Stevens, E. Falconer, J. A. Goldsmith, C.-W. Chou, O. M. Abiona, A. West, K. Westendorf, K. Muthuraman, E. J. Fritch, K. H. Dinnon, A. Schäfer, M. R. Denison, J. D. Chappell, R. S. Baric, B. S. Graham, K. S. Corbett, J. S. McLellan, Stabilized coronavirus spike stem elicits a broadly protective antibody. Cell Reports 37, 109929 (2021).

75. W. Pang, Y. Lu, Y.-B. Zhao, F. Shen, C.-F. Fan, Q. Wang, W.-Q. He, X.-Y. He, Z.-K. Li, T.-T. Chen, C.-X. Yang, Y.-Z. Li, S.-X. Xiao, Z.-J. Zhao, X.-S. Huang, R.-H. Luo, L.-M. Yang, M. Zhang, X.-Q. Dong, M.-H. Li, X.-L. Feng, Q.-C. Zhou, W. Qu, S. Jiang, S. Ouyang, Y.-T. Zheng, A variant-proof SARS-CoV-2 vaccine targeting HR1 domain in S2 subunit of spike protein. Cell Res 32, 1068– 1085 (2022).

76. P. J. Halfmann, S. J. Frey, K. Loeffler, M. Kuroda, T. Maemura, T. Armbrust, J. E. Yang, Y. J. Hou, R. Baric, E. R. Wright, Y. Kawaoka, R. S. Kane, Multivalent S2-based vaccines provide broad protection against SARS-CoV-2 variants of concern and pangolin coronaviruses. eBioMedicine 86, 104341 (2022).

77. P. Zhou, G. Song, H. Liu, M. Yuan, W. He, N. Beutler, X. Zhu, L. V. Tse, D. R. Martinez, A. Schäfer, F. Anzanello, P. Yong, L. Peng, K. Dueker, R. Musharrafieh, S. Callaghan, T. Capozzola, O. Limbo, M. Parren, E. Garcia, S. A. Rawlings, D. M. Smith, D. Nemazee, J. G. Jardine, Y. Safonova, B. Briney, T. F. Rogers, I. A. Wilson, R. S. Baric, L. E. Gralinski, D. R. Burton, R. Andrabi, Broadly neutralizing anti-S2 antibodies protect against all three human betacoronaviruses that cause deadly disease. Immunity 56, 669–686.e7 (2023).

78. K. W. Ng, N. Faulkner, K. Finsterbusch, M. Wu, R. Harvey, S. Hussain, M. Greco, Y. Liu, S. Kjaer, C. Swanton, S. Gandhi, R. Beale, S. J. Gamblin, P. Cherepanov, J. McCauley, R. Daniels, M. Howell, H. Arase, A. Wack, D. L. V. Bauer, G. Kassiotis, SARS-CoV-2 S2–targeted vaccination elicits broadly neutralizing antibodies. Sci. Transl. Med. 14, eabn3715 (2022).

79. F. A. Lempp, L. B. Soriaga, M. Montiel-Ruiz, F. Benigni, J. Noack, Y.-J. Park, S. Bianchi, A. C. Walls, J. E. Bowen, J. Zhou, H. Kaiser, A. Joshi, M. Agostini, M. Meury, E. Dellota, S. Jaconi, E. Cameroni, J. Martinez-Picado, J. Vergara-Alert, N. Izquierdo-Useros, H. W. Virgin, A. Lanzavecchia, D. Veesler, L. A. Purcell, A. Telenti, D. Corti, Lectins enhance SARS-CoV-2 infection and influence neutralizing antibodies. Nature 598, 342–347 (2021).

80. J. E. Bowen, A. Addetia, H. V. Dang, C. Stewart, J. T. Brown, W. K. Sharkey, K. R. Sprouse, A. C. Walls, I. G. Mazzitelli, J. K. Logue, N. M. Franko, N. Czudnochowski, A. E. Powell, E. Dellota, K. Ahmed, A. S. Ansari, E. Cameroni, A. Gori, A. Bandera, C. M. Posavad, J. M. Dan, Z. Zhang, D. Weiskopf, A. Sette, S. Crotty, N. T. Iqbal, D. Corti, J. Geffner, G. Snell, R. Grifantini, H. Y. Chu, D. Veesler, Omicron spike function and neutralizing activity elicited by a comprehensive panel of vaccines. Science 377, 890–894 (2022).

81. A. Addetia, L. Piccoli, J. B. Case, Y.-J. Park, M. Beltramello, B. Guarino, H. Dang, G. D. De Melo, D. Pinto, K. Sprouse, S. M. Scheaffer, J. Bassi, C. Silacci-Fregni, F. Muoio, M. Dini, L. Vincenzetti, R. Acosta, D. Johnson, S. Subramanian, C. Saliba, M. Giurdanella, G. Lombardo, G. Leoni, K. Culap, C. McAlister, A. Rajesh, E. Dellota, J. Zhou, N. Farhat, D. Bohan, J. Noack, A. Chen, F. A. Lempp, J. Quispe, L. Kergoat, F. Larrous, E. Cameroni, B. Whitener, O. Giannini, P. Cippà, A. Ceschi, P. Ferrari, A. Franzetti-Pellanda, M. Biggiogero, C. Garzoni, S. Zappi, L. Bernasconi, M. J. Kim, L. E. Rosen, G. Schnell, N. Czudnochowski, F. Benigni, N. Franko, J. K. Logue, C. Yoshiyama, C. Stewart, H. Chu, H. Bourhy, M. A. Schmid, L. A. Purcell, G. Snell, A. Lanzavecchia, M. S. Diamond, D. Corti, D. Veesler, Neutralization, effector function and immune imprinting of Omicron variants. Nature 621, 592–601 (2023).

82. A. Addetia, Y.-J. Park, T. Starr, A. J. Greaney, K. R. Sprouse, J. E. Bowen, S. W. Tiles, W. C. Van Voorhis, J. D. Bloom, D. Corti, A. C. Walls, D. Veesler, Structural changes in the SARS-CoV-2 spike E406W mutant escaping a clinical monoclonal antibody cocktail. Cell Rep 42, 112621 (2023).

83. Q. Xiong, L. Cao, C. Ma, M. A. Tortorici, C. Liu, J. Si, P. Liu, M. Gu, A. C. Walls, C. Wang, L. Shi, F. Tong, M. Huang, J. Li, C. Zhao, C. Shen, Y. Chen, H. Zhao, K. Lan, D. Corti, D. Veesler, X. Wang, H. Yan, Close relatives of MERS-CoV in bats use ACE2 as their functional receptors. Nature 612, 748–757 (2022).

